# A multiomic atlas of human early skeletal development

**DOI:** 10.1101/2024.07.10.602965

**Authors:** Ken To, Lijiang Fei, J. Patrick Pett, Kenny Roberts, Krzysztof Polański, Tong Li, Nadav Yayon, Peng He, Chuan Xu, James Cranley, Ruoyan Li, Kazumasa Kanemaru, Ni Huang, Stathis Megas, Laura Richardson, Rakesh Kapuge, Shani Perera, Elizabeth Tuck, Anna Wilbrey-Clark, Ilaria Mulas, Fani Memi, Batuhan Cakir, Alexander V. Predeus, David Horsfall, Simon Murray, Martin Prete, Pavel Mazin, Xiaoling He, Kerstin B. Meyer, Muzlifah Haniffa, Roger A. Barker, Omer Bayraktar, Christopher D. Buckley, Sarah A. Teichmann

## Abstract

Bone and joint formation in the developing skeleton rely on co-ordinated differentiation of progenitors in the nascent developing limbs and joints. The cell states, epigenetic processes and key regulatory factors underlying their lineage commitment to osteogenic and other mesenchymal populations during ossification and joint formation remain poorly understood and are largely unexplored in human studies. Here, we apply paired single-nuclei transcriptional and epigenetic profiling of 336,000 droplets, in addition to spatial transcriptomics, to construct a comprehensive atlas of human bone, cartilage and joint development in the shoulder, hip, knee and cranium from 5 to 11 post-conception weeks. Spatial mapping of cell clusters to our highly multiplexed in situ sequencing (ISS) data using our newly developed tool ISS-Patcher revealed new cellular mechanisms of zonation during bone and joint formation. Combined modelling of chromatin accessibility and RNA expression allowed the identification of the transcriptional and epigenetic regulatory landscapes that drive differentiation of mesenchymal lineages including osteogenic and chondrogenic lineages, and novel chondrocyte cell states. In particular, we define regionally distinct limb and cranial osteoprogenitor populations and trajectories across the fetal skeleton and characterise differential regulatory networks that govern intramembranous and endochondral ossification. We also introduce SNP2Cell, a tool to link cell-type specific regulatory networks to numerous polygenic traits such as osteoarthritis. We also conduct *in silico* perturbations of genes that cause monogenic craniosynostosis and implicate potential pathogenic cell states and disease mechanisms involved. This work forms a detailed and dynamic regulatory atlas of human fetal skeletal maturation and advances our fundamental understanding of cell fate determination in human skeletal development.

## Main

Human bone formation begins between 6-8 post-conception weeks (PCW) in the period spanning the transition from embryonic to fetal stages. Within the calvaria of the cranial skeleton, progenitors differentiate into osteoblasts through intramembranous (IM) ossification and expand from ossification centres that eventually meet to establish the prenatal suture joints that house osteoprogenitors ^1,2^. In the appendicular skeleton, the nascent synovial joint first appears as an interzone condensation in the limb bud at 5-6 PCW ^3^, which subsequently forms a cavitated region that articulates adjacent incipient cartilage templates. The latter acts as a temporary scaffold to facilitate development of the body plane and is gradually replaced by bone tissue as development proceeds through endochondral (EC) ossification ^4,5^.

These two distinct and anatomically restricted modes of ossification govern osteogenesis and joint formation throughout the human skeleton and, to our knowledge, the cellular bases by which they form and mature remain partially described in human development at single-cell resolution. To address this, we applied single-nuclei paired RNA and ATAC sequencing, and multiple spatial methods to decipher the gene regulatory networks that mediate maturation of the distinct bone and joint-forming niches in the cranium and appendicular skeleton across space and time from 5-11 PCW. Through this, we discovered previously undescribed cellular diversity in the osteogenic and chondrogenic lineages. We develop ISS-Patcher, a tool to impute cell labels from the droplet data on our high-resolution 155-plex in situ sequencing (ISS) datasets, allowing us to gain detailed insights into spatially defined niches within the embryonic synovial joint. Applying OrganAxis, a new spatial transcriptomics annotation tool, we also define the spatial trajectory of the developing frontal bone of the fetal skull.

In addition, our comprehensive resource and new computational toolset including SNP2Cell enabled us to gain new insights into the molecular mechanisms of developmental bone disease, such as craniosynostosis^6–8^ as well as to implicate region-specific contributions of bone and cartilage lineages in ageing diseases of the human skeleton, such as osteoarthritis^9,10^.

## Results

### Cellular taxonomy across first trimester skeletal joint development

We applied paired droplet-based snRNA-seq and snATAC-seq (10x Genomics Multiome) to define the cellular taxonomy of the developing embryonic fetal skeleton between 5-11 post conception weeks (PCW) across 12 fetal donors. Focusing on three major synovial joints, we sampled whole-intact nascent and maturing shoulder, hip and knee joints across the developmental timepoints (Fig. 1a-b). For the developing skull and cranial suture joints, which had not been profiled in human across embryonic stages, we studied the anterior and posterior regions of the cranium separately, and sampled the calvaria and skull base individually to divide intramembranous and endochondral bone-forming niches. (Fig. 1a and Supplementary Table 1). We captured 336,162 high-quality transcriptome and chromatin-accessibility droplets and curated the dataset from all regions into eight shared major cellular compartments based on marker gene expression: mesenchymal, muscle, immune, endothelial, Schwann, neural, epithelial, and erythroid cells (Fig. 1c-e, Extended Data Fig. 1c). High concordance was observed between the transcriptome and ATAC peak compartment structures (Extended Data Fig. 1f), and we observed that mesenchymal cells were the most abundant across all regions, while myogenic cells were absent in the calvaria (Fig. 1e). From these we defined 30 broad cell clusters and conducted sub-clustering of lineages which enabled the identification of 122 fine-grained clusters (Supplementary Table 2).

**Figure. 1:**
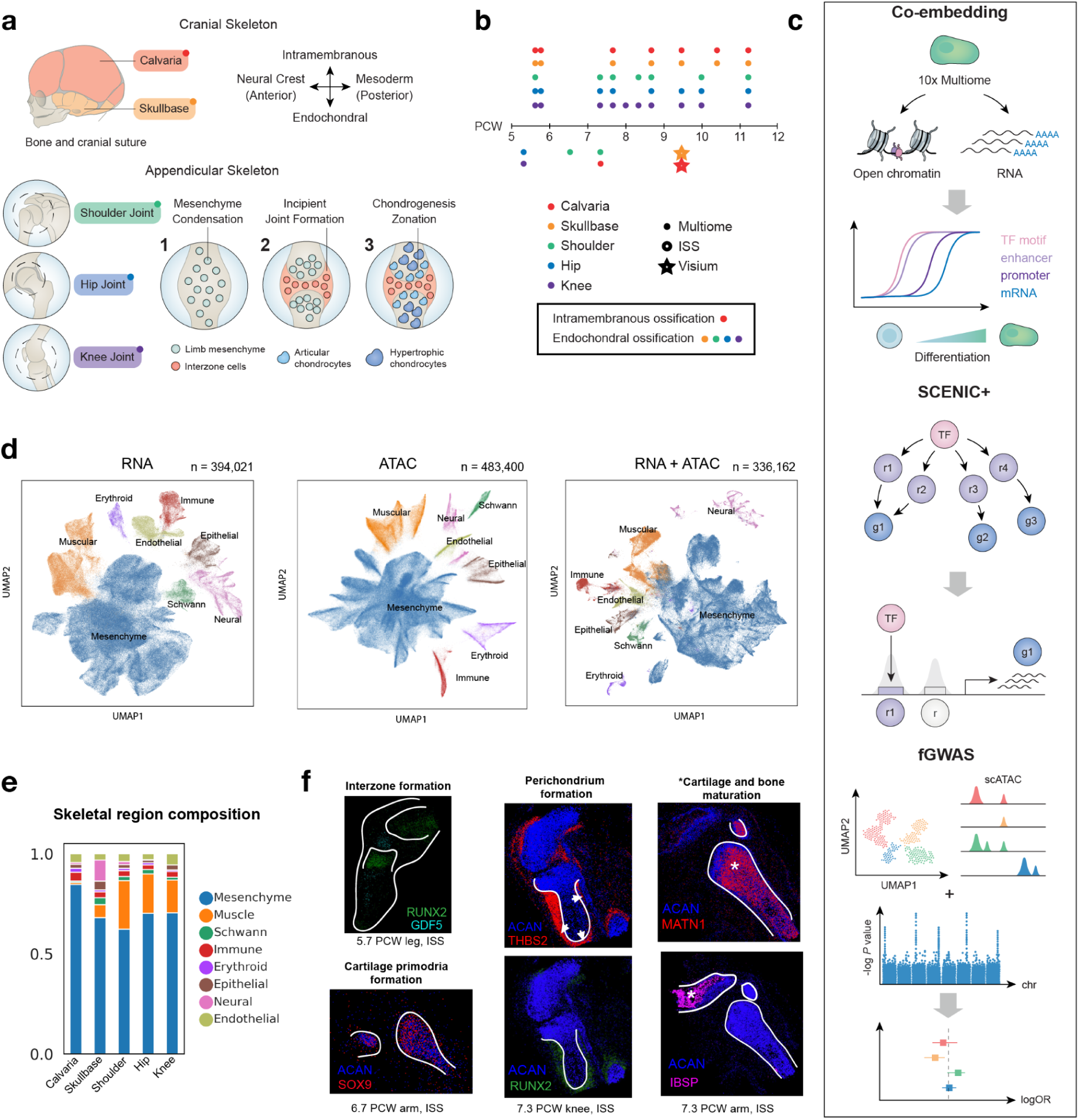
Human fetal skeletal atlas overview. **a)** Anatomical sampling approach to dissect five main anatomical sites. Cell types of origin within the cranium are determined by the anterior-posterior axis. **b)** Donor overview across age and anatomical regions sampled; atlasing modalities represented in the legend. **c)** Analysis approaches applied to integrated RNA-ATAC seq data. **d)** UMAP embedding of dataset using RNA-seq only, ATAC-seq only and integrated datasets. **e)** Cellular compartment composition of the skeletal region across anatomical regions. **f)** Relative cell type abundance across anatomical locations.

Previous single-cell (scRNA-seq) atlases of developing human and mouse limbs ^11–13^ profiled whole cells and captured low numbers of maturing osteoblast transcriptomes and *COL10A1*+ hypertrophic chondrocyte transcriptomes. To facilitate the reconstruction of developmental trajectories in osteochondral lineages, we profiled nuclei-droplets from a large number of cells. This captured comparatively more diverse chondrogenic and osteogenic subcompartments, suggesting that matrix-rich stromal populations are relatively resistant to enzymatic digestion and less amenable to whole-cell profiling ^11–13^.

Our approach enabled the discovery of osteogenic cell state trajectories enriched in the appendicular joints and skull base (formed through endochondral ossification) and calvarium (formed through intramembranous ossification), reflecting different osteoblast origins from anatomically distinct regions (Extended Data Fig. 2). Chondrogenic clusters were relatively depleted in the calvarium (Fig. 2a), consistent with the mechanisms of intramembranous bone formation^14^. In addition, we uncovered novel cell states of the facial and pharyngeal regions, described *COL10A1*-expressing fetal hypertrophic chondrocyte development, discovered a novel *PAX7*^+^ chondrocyte population, and described a potential pathway leading to its formation from common myogenic progenitors^15^. We delineate the previously unreported process of human Schwann cell development and the formation of fetal *PI16*^+^ fibroblasts, the latter of which shares transcriptional similarities with postnatal pan-tissue perivascular fibroblasts.

**Figure. 2:**
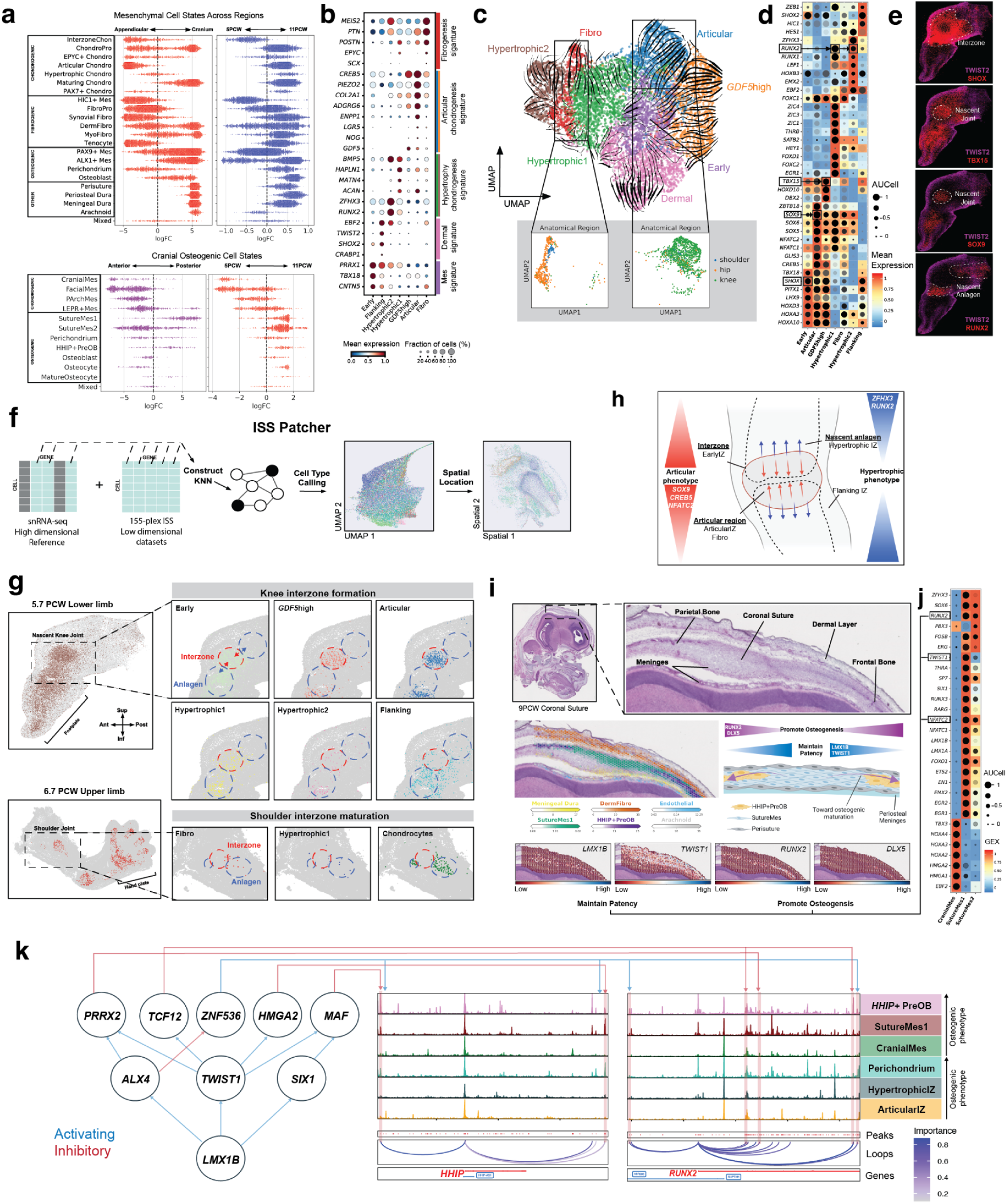
Formation of the embryonic joints across space and time. **a)** Differential abundance (MILO) of cell states across lineages against multiple comparators. Top: Broad clusters of non-myogenic mesenchyme. Bottom: Cranial osteogenic subclusters. ChondroPro: Chondroprogenitors, *EPYC*^+^ Chondro: *EPYC*^+^ Chondrocyte, Articular Chondro: Chondrocyte, Hypertrophic Chondro: Hypertrophic Chondrocyte, Maturing Chondro: Maturing Chondrocyte, *PAX7*^+^ Chondro: *PAX7*^+^ Chondrocyte, *HIC1*^+^ Mes: *HIC1*^+^ Mesenchyme, FibroPro: Fibroblast progenitors, Synovial Fibro: Synovial fibroblast, DermFibro: Dermal fibroblast, MyoFibro: Myofibroblast, *PAX9*^+^ Mes: *PAX9*^+^ Mesenchyme, *ALX1*^+^ Mes, *ALX1*^+^ Mesenchyme, CranialMes: Cranial mesenchyme, FacialMes: Facial mesenchyme, PArchMes: Pharyngeal mesenchyme, *LEPR*^+^ Mes: *LEPR*^+^ Mesenchyme, SutureMes1: Suture mesenchyme1, SutureMes2: Suture mesenchyme2, *HHIP*^+^ PreOB: *HHIP*^+^ Pre-Osteoblast. Mixed: other related cell types that are not included in each plot respectively. **b)** Normalised gene expression of genes in modules associated with skeletal development within InterzoneChon subclusters. **c)** RNA-velocity on UMAP of RNA-sub-clustered cell states of the broad InterzoneChon cluster. **d)** SCENIC+ predicted TF expression (box color), dot size shows target gene accessibility (AUCell) and dot shade (grayscale) shows target gene expression (GEX AUCell). **e)** Demultiplexed marker gene expression from ISS of the 5.7 PCW lower limb. **f)** ISS-Patcher workflow (see methods). **g)** Spatial plots of ISS-Patcher cell cluster imputation. **h)** Schematic of knee joint interzone formation, gradients demonstrating genes associated with zonated hypertrophic and articular phenotypes. **i)** Histological view and annotations of adjacent sections of 10x cytassist visium data. Cell2location results of coronal suture and schematic demonstrating TF gradient across regions of the coronal suture. **j)** SCENIC+ predicted TF expression (box color), dot size shows target gene accessibility (AUCell) and dot shade (grayscale) shows target gene expression (GEX AUCell) in suture progenitors. **k)** Coverage plots showing aggregated scATAC signals around the *HHIP* and *RUNX2* loci for osteoprogenitor cell states with increasing osteogenic phenotype in IM (CranialMes to HHIP+ ProOB) and EC (ArticularIZ to Perichondrium). Below each coverage plot, loops predicted by Scenic+ between TSS and enhancers are shown (colored by importance score). Selected upstream TFs predicted by Scenic+ to bind and regulate via some of the enhancers are shown on the left. The network links inhibitors of osteogenesis (such as TWIST1, LMX1B) to pro-osteogenic genes (RUNX2, HHIP) via overall inhibitory connections.

Sequencing-based spatial transcriptomics previously deployed on the embryonic limb has highlighted challenges in capturing transcripts within the early ossifying bone and cartilaginous precursors (cartilage anlagen)^11^. In order to resolve bone lineage cell states in space and understand organisation of the nascent synovial joint that bridges the adjacent anlagen, we performed high resolution 155-plex ISS of the whole intact early embryonic upper (6.7 PCW) and lower (5.7 PCW) limbs, late embryonic (7.3 PCW) knee, and shoulder regions (Fig. 1f and Extended Data Fig. 2a-c). Additionally, we conduct sequencing-based spatial transcriptomics (10x Genomics Visium CytAssist) of the developing coronal suture (9 PCW) and frontal bone (Extended Data Fig. 2d) allowing us to capture the trajectory of osteo-lineage development across space. We leveraged these spatial data to systemically curate cell lineages within the mesenchymal compartment in a well-defined spatial context.

### Spatiotemporal zonation of the incipient synovial joints

Synovial joint-site determination occurs between 5-6 PCW in the limbs, orchestrated by the emergence of a mesenchymal condensation comprising *GDF5*-expressing populations termed the interzone (IZ). Part of them undergo differentiation into joint-articulating chondrocytes at the ends of the endochondral bone primordia, and chondrocytes that adopt a hypertrophic phenotype in the cartilage anlagen^16^. To identify cell-states that constitute these early joint-specific progenitors, we applied differential abundance testing focusing on the non-myogenic mesenchyme across developmental time (5-11 PCW) and skeletal regions (shoulder, hip, knee) (Fig. 2a). We identified four progenitor populations enriched in the early appendicular joint: Interzone Chondrocytes (InterzoneChon), *HIC1*^+^ mesenchyme (*HIC1*^+^ Mes), PI16-expressing fibroblast-progenitors (FibroPro) and dermal fibroblasts (DermFibro). These clusters support the notion that numerous early progenitors exist between 5-7 PCW and give rise to the chondro-fibro lineages in the joint, instead of a single master *GDF5*^+^ progenitor population^17^.

We first focused on deciphering the composition of progenitors in the early synovial joint, where fibrous ligaments^18^, tendon^19^ and cartilage components are thought to derive from a progenitor *GDF5*^+^ population in mice^17,20^. To this end, we performed sub-clustering of the broad InterzoneChon (*GDF5*^+^) population and leveraged RNA-velocity to infer their pseudo-trajectory (Fig. 2b-c). We then applied SCENIC+ to identify predicted gene programs and TF accessibility changes across the resulting subclusters (Fig. 2d). Our analysis revealed multiple subclusters with specific tissue expression signatures, including an early *PRRX1*^+^ mesenchyme population (EarlyIZ) enriched for mesenchyme-associated signatures (Fig. 2b). Early IZ was predicted to express TFs associated with early limb development (e.g. *TBX18*, *SHOX*, *LHX9*)^11^ and demonstrated low *RUNX2* expression and accessibility in downstream target regions but moderate *SOX5, SOX6* and *SOX9* expression and target region accessibility, suggesting a poised trajectory favouring chondrogenesis over osteogenesis (Fig. 2d).

We defined four main trajectories that emerge from this latter population, characterised by transcriptional signatures of fibroblasts (Fibro IZ), articular (Articular IZ) and hypertrophic chondrocytes (Hypertrophic IZ-1, Hypertrophic IZ-2), and flanking dermal mesenchyme (Flanking IZ) (Fig. 2b-c). Three clusters; Articular, Fibro and *GDF5^high^* IZ, expressed *GDF5*. Articular IZ was more prevalent in the knee joint, whereas Fibro IZ is enriched in the ball-and-socket joints of the shoulder and hip (Fig. 2c). Articular IZ particularly expressed chondrogenic markers (*COL2A1*, *ADGRG6*, *PIEZO2*, *LGR5*, *NOG)* comparable to mouse IZ progenitors for articular chondrocytes ^17,20^ including *ENPP1*, which negatively regulates chondrocyte hypertrophy and bone formation, in keeping with a role in articular cartilage formation. While *GDF5*-high IZ was predicted to highly express *CREB5* and *DBX2* and its target genes, the latter of which is associated with digital IZ formation regulated by *HOX* genes^17,21^, chondrogenesis TFs were not highly expressed, and predicted TF activity and accessibility for *RUNX2* was lowest among the IZ populations. We therefore hypothesise that *GDF5high* IZ is maintained as a progenitor pool, potentially to sustain continuous influx into the forming joint^18^. In the *GDF5* negative hypertrophic clusters, progressive TF expression and accessibility of *RUNX2*^22–24^, and concurrent downregulation of articular chondrocyte TFs (*CREB5*, *EGR1*) signifies a hypertrophic phenotype (Fig. 2d). Hypertrophic2 IZ transitions through Hypertrophic1 in the inferred trajectory and the former expressed genes that are upregulated during formation of anlagen or pre-hypertrophic chondrocytes in mice (*BMP5*, *HAPLN1*, *ACAN* and *ZFHX3*)^19,25^, suggesting they form the incipient cartilage template. Overall, our *in vivo* data are consistent with *in vitro* observations of the propensity for human-iPSC and mouse-ESC derived *GDF5*^+^ stromal cells to form an articular, rather than a hypertrophic phenotype ^26,27^.

To understand spatial zonation of the nascent synovial joint we next visualised the pre-cavitating knee joint ISS to identify regions of incipient hypertrophic anlagen (*RUNX2*), early progenitors (*TBX15*, *SHOX*) and nascent articular cartilage and interzone (*SOX9*) (Fig. 2e). We then leveraged the snRNA-seq dataset and imputed genes missing in the 155-plex ISS dataset through our newly developed ISS-Patcher function (see Methods) to infer cell labels to the 155-plex clustered manifold (Fig. 2f,g). In the embryonic lower limb, Early IZ was diffusely distributed across regions of the interzone and anlagen, and was surrounded by Flanking IZ (Fig. 2c,d). Articular IZ (*GDF5*^+^), were predominantly enriched in sites of incipient knee articular cartilage formation which also showed SOX9 staining (Fig. 2e,g). In contrast, Fibro IZ (*GDF5*^+^) was enriched in the shoulder interzone region adjacent to the articular surface of the humerus (Fig. 2g), and expressed *SCX* (Fig. 2b) which plays a role in ligament formation in the IZ in mice^28^. The differential enrichment and signatures of the Articular and Fibro IZ populations may be explained by the need for fibro-cartilage labrum formation in the ball-and-socket joints of the shoulder and hip which emerge and mature later in development. We coalesce the trajectories and spatial enrichment of the newly defined interzone progenitors and describe a model for zonation of the embryonic joint. Within the model, the future joint region (Articular phenotype) forms in the centre of the interzone defined by early pro-chondrogenic and anti-osteogenic TF enrichment, the incipient fetal anlagen (Hypertrophic phenotype) forms at the edges away from the joint, defined by *RUNX2* enrichment (Fig. 2h).

### Emergence of fibroblast lineages in synovial joints

Fibroblast lineage cell states have previously been described in the developing mouse limb to arise from a master *HIC1*^+^ precursor population^29^, which contributes minimally to osteochondral components. In addition, a “universal” *PI16*^+^ population is thought to persist in postnatal stages and govern adult fibroblast formation across tissues in mice and humans ^29,30^. Here, we sought to uncover the taxonomy of the fibroblast lineage in first trimester human joints. We first identified fibroblast progenitors (FibroPro) and *HIC1*^+^ mesenchyme (*HIC1*^+^ Mes) enriched in the appendicular joints during the embryonic period (<8 PCW), surrounding the nascent joint and with diffuse distribution in the limbs, respectively (Fig. 2a, Extended Data Fig.3a-c,f). *TWIST1*, a known activator of postnatal fibrosis and *TWIST2* a regulator of postnatal dermal fibroblast proliferation ^31,32^, were predicted to show high TF activity in DermFibro (Extended Data Fig. 3e), which was present across sites and time, and localised to the skin around the developing embryonic limbs (Extended Data Fig.3 f-g). Interestingly, *EN1*, a TF required for the fibrotic response and associated scarring during postnatal wound healing^33^, was highly expressed in DermFibro (Extended Data Fig. 3e) but showed low target gene expression, suggesting target repression, consistent with observations of scarless wound healing in utero. In the cranium, it formed the majority of early-stage enriched fibro-lineage progenitors (Fig. 2a).

To uncover developmental dynamics in the appendicular joints, we reconstructed pseudotime trajectory across fibroblasts and predicted *HIC1*^+^ Mes as a progenitor to FibroPro during the embryonic phase at <8 PCW (Extended Data Fig. 3a). *HIC1*^+^ Mes enriched for activity in numerous proliferation associated TFs including *WT1*, *SOX5*, and *FOXC1*, which is associated with an invasive and activated synovial fibroblast phenotype (Extended Data Fig. 3e) ^34–37^. At ∼8 PCW, FibroPro forms a “fibroblast hub” and expresses *PI16* and *DPT*, markers of pan-tissue adventitia-associated fibroblasts in postnatal health (Extended Data Fig. 3d). Here, it gives rise to tenocytes, synovial, dermal and myo- fibroblasts (Fig. 2a,b). *BNC2*, a myofibroblast-associated TF ^38^, had high activity in FibroPro, consistent with its postnatal pan-tissue presence in the vascular adventitia ^30^. Additionally, *YBX1*, a TF shown to drive proliferation of mouse embryonic fibroblasts was also enriched. *HIC1*+ Mes additionally differentiated to the tenogenic lineage during the embryonic, and synovial fibroblasts in the fetal phase, highlighting potential parallel routes to synovial fibroblast formation in the prenatal skeleton (Extended Data Fig. 3a-d).

### Formation of the cranium and suture joints

In the cranium, incipient suture mesenchyme matures beyond 7 PCW at meeting points of bone fronts emanating from primary ossification centres in the flat bones, forming suture joints^39^. We conducted differential abundance testing of the mesenchyme compartment and revealed the embryonic cranium (<7 PCW) was dominated by dermal and myo- fibroblasts, and two broad *PAX9*^+^*RUNX2*^+^ and *ALX1*^+^*RUNX2*^+^ mesenchymal clusters that had not been described previously in human craniogenesis (Fig. 2a). The latter expressed markers such as *TWIST1*, *CTSK*, *ZIC1*, consistent with mouse cranial progenitors^40^, and *RUNX2*, suggesting they form part of the osteogenic lineage (Extended Data Fig. 4a). *ALX1* is required for cranium formation in the mouse^41^, and neural crest cell (NCC) migration and differentiation in human derived stem cells, with variants of the gene associated with frontonasal dysplasia ^28,42^. Single-cell studies of the human brain have also identified *ALX1* expression within mesenchyme-like progenitors^43^. We therefore used metadata obtained during tissue sampling to examine differences in anterior-posterior skull gene expression and found numerous neural crest cell (NCC) TFs and targets linked in our GRN including *ALX1*, *PAX3*, *BMP5*, *COL4A2*, *COL4A1* and *TSHZ2* which are differentially enriched in the anterior portion of the cranium (Extended Data Fig. 4b). This suggests that *ALX1*^+^ Mes may have a neural crest origin. Due to the transient embryonic nature of NCCs^44^ we did not capture early embryonic bona fide *SOX10*^+^ NCCs. *RUNX2*-expressing *PAX9*^+^ Mes was present in the cranium (skull base) and appendicular skeleton (Fig. 2a), and therefore likely represents a lateral plate and axial mesoderm-derived progenitor for osteoblasts in endochondral ossification.

Next, we sought to delineate the contribution of the broad *ALX1*^+^ and *PAX9*^+^ Mes clusters to the bone-forming suture joint. We first sub clustered these populations in combination with osteoblasts (*SP7*, *ALP*), osteocytes (*SOST*, *DMP1*) and perichondrium (*RUNX2*, *THBS2*, *POSTN*) to form an osteo-lineage subcompartment. This revealed anatomical and age -segregated clusters (Fig. 2a, Extended Data Fig. 4c-d). Among these were three early progenitors, including facial (FacialMes) and pharyngeal mesenchyme (PArchMes) which were located in the skull base and expressed markers of axial mesoderm (*PAX3* and *LHX8*), and a cranial mesenchyme (CranialMes) population which was abundant in the calvarium (Fig. 2a, Extended Data Fig. 4d-e).

Cranial sutures form in fetal stages following expansion of the primary ossification centres at the end of the embryonic period^1^. Consistent with this, we define two human suture mesenchyme clusters (SutureMes1/2) enriched in the cranium from 8 PCW (Fig. 2a). Classical markers of fetal cranial sutures (*TWIST1*, *ZIC1*, *ZIC4*), previously reported in NCC-derived mesenchyme in mice, were enriched in both populations in addition to high expression levels of *THBS2*, akin to the endochondral perichondrium cluster (Extended Data Fig. 4d)^45^. Notably, the SutureMes populations expressed *CTSK*, a marker of periosteal mesenchymal stem cells in the postnatal mouse cranial osteogenic niche shown to mediate intramembranous ossification^46^. Using Cell2location, we revealed the spatial distribution of SutureMes1 and SutureMes2 within the developing coronal suture joint, spanning outward toward the periosteum (Fig. 2i). Osteoprogenitors (*HHIP*^+^PreOB) emerged at the opposing frontal and parietal bone boundaries of the suture populations (Fig. 2i). Additionally, the sutures were flanked by peri-suture (*CTSK*^-^) in contrast to the *CTSK*^+^ postnatal mouse periosteum previously reported^46^ (Extended Data Fig. 4g).

Analogous to osteogenic repressor genes enriched in spatially-defined clusters of the articular IZ, SutureMes populations also showed predicted TF and target accessibility enrichment of negative regulators of osteogenesis in the mouse cranial sutures (*TWIST1*, *LMX1B*, *NFATC2*)^47^ (Fig. 2i). Numerous TFs associated with osteogenesis (*EGR1*, *EGR2*, *SP7*)^48,49^ and osteoblast formation (*FOXO1*, *ETS1*, *ETS2*)^50^ were predicted suggesting a transcriptional network primed for impending osteogenesis. Akin to the molecular gradients of the joint-interface in the embryonic knee (Fig. 2h), we observed *LMX1B* and *TWIST1* expression within the suture region, dissipating toward the flanking bone edges alongside a rise in regulators of osteogenesis such as *RUNX2* (Fig. 2i), suggesting comparable mechanisms in sustaining the non-osteogenic interface in both the limb and skull bones.

To dissect this process and uncover the enhancer-driven gene regulatory network (eGRN) of the loci surrounding key osteogenic transcription factors *RUNX2* and calvaria-enriched *HHIP*, we visualised coverage of the network predicted by SCENIC+. Both *RUNX2* and *HHIP* were predicted to be inhibited by a shared set of anti-osteogenic TFs including *LMX1B*, *TWIST1* and ALX4 via several intermediate repressors targeting various enhancers around their loci (Fig. 2k), illustrating the relationships maintaining the balance of osteogenesis initiation. *HHIP* was most accessible in *HHIP*^+^PreOB and was indirectly repressed by *LMX1B* via *TWIST1* which positively regulated its immediate repressors *MAF* and *HMGA2* via two differentially accessible enhancers of *HHIP* enriched in SutureMes1 and *HHIP*+PreOB. *RUNX2* was most accessible in *HHIP*^+^PreOB (IM) as well as Perichondrium (EC) and was predicted to be indirectly targeted by the same repressors via *TCF12* and *PRRX2*. Overall, the network illustrates the coherent regulation of bone-adjacent non-ossifying niches by key osteogenic regulators via multiple redundant paths.

### Distinct trajectories of osteogenesis across the skeleton

Osteoblastogenesis, signifying the onset of ossification, commences from ∼7-8 PCW onward (Fig. 2a). To reconstruct this process, we utilised force-directed embedding on non-cycling cells and uncovered two major trajectories from distinct progenitors converging toward osteoblast and osteocyte formation (Fig. 3a). Endochondral ossification (EC) consisted of two routes, stemming from limb mesenchyme (LimbMes), FacialMes and PArchMes. LimbMes, a cluster which encompasses upper and lower limb joints, demonstrated transcriptional similarity (*ISL1*, *TBX5*, *WT1*) to lateral plate mesoderm previously reported in the fetal human limb bud (Fig. 3a, Extended Data Fig. 4a-e) ^11,30^. CranialMes and FacialMes were differentially abundant in the anterior portion of the calvarium and skull base (Fig. 2a), respectively. FacialMes expressed NCC-derived mesenchyme regulators *PAX3* and *PAX7* (Extended Data Fig. 4e)^51^. Likewise, *ALX1* was enriched in CranialMes, leading us to hypothesise that the clusters represent, to our knowledge, previously undescribed human NCC-derived osteogenic populations. *LEPR*+Mes was enriched in the calvarium, *LEPR* has been reported as a marker of second trimester and post-natal osteogenic stromal population ^12,52^. PArchMes emerged between 6-8 PCW and expressed marker genes of the first pharyngeal arch (*PAX9*, *LHX*, *DLK1*) ^53^, and is likely to comprise mixed origins from axial mesoderm and NCC. We discovered a two-stage process in the intramembranous pathway (IM), defined by the transition between embryonic and fetal stages. Suturogenesis occurs in stage 1, as CranialMes differentiates into SutureMes1/2 (Fig. 3a). In stage 2, an osteogenic trajectory emerges from SutureMes1 toward SutureMes2, followed by *HHIP*^+^PreOB. *HHIP* was previously reported in the mouse as a marker of osteogenic coronal suture mesenchyme ^54^. We demonstrate here that they are a distinct population progeny to *TWIST1*-enriched SutureMes and are distributed at the boundaries of bone formation in human fetal development (Fig. 2i).

**Figure 3:**
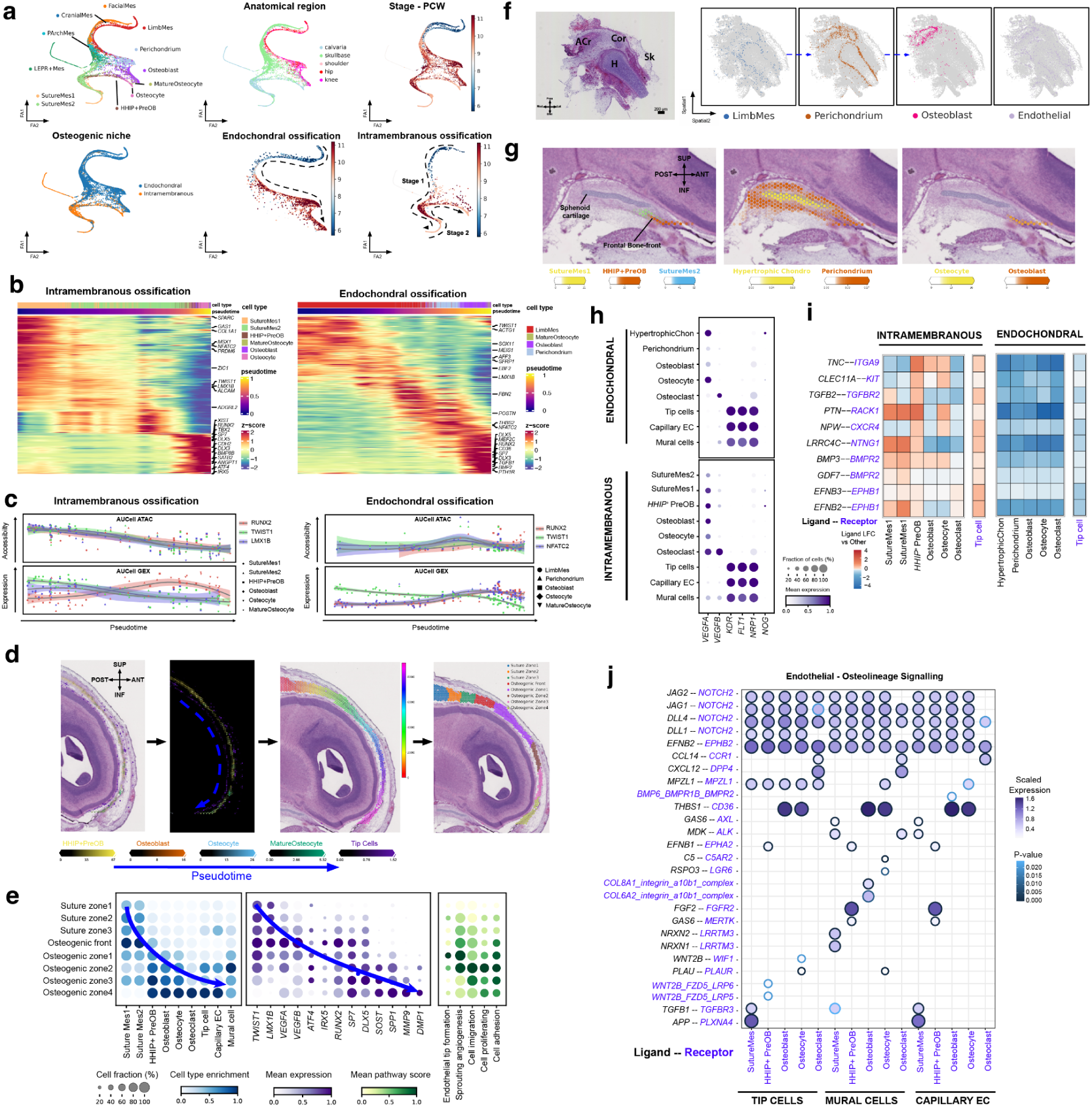
Intramembranous and endochondral osteogenesis. **a)** Force directed (FA) embedding showing trajectories per anatomical region that converge on osteoblasts and osteocytes. In intramembranous (IM) ossification the suture mesenchyme constitutes a distinct state that maintains progenitor pools. A two-stage IM process forms the sutures then subsequently bone-lineage. LimbMes: Limb mesenchyme. **b)** Heatmaps showing the expression of differentially expressed genes along pseudotime for IM and endochondral (EC) ossification. Differential expression has been tested using a spatial autocorrelation test in monocle3. **c)** TF accessibility (AUCell ATAC) and Target gene accessibility (AUCell GEX) of selected TFs along pseudotime. Differing patterns are observed for osteogenic (RUNX2) and inhibitory TFs (TWIST1, LMX1B, NFATC2), pointing to a complex regulatory mechanism. **d)** Spatial binning of Visium Cytassist image of frontal bone (sagittal section) using OrganAxis, axis begins from coronal suture to the ossifying frontal bone (posterior-anterior) Left-Right: Histological image of frontal bone; Cell-cluster enrichment from Cell2location; Axis values (rainbow); manual bins based on histological features **e)** Enrichment within spatial bins for IM cell states (left), selected marker gene expression (middle) and pathway enrichment (right). **f)** 7.3 PCW humerus (Coronal section) and imputed cell clusters showing EC ossification as a sequence of mapped cell-states. ACr, Acromion; Cor, Coracoid; H, Humerus; Sk, Skin. **g)** Cross-section of Visium Cytassist at the meeting point between calvaria and skull base (anterior border of the sphenoid), chondrogenic cell states enrichment from Cell2location **h)** Predicted ligand-receptor interactions per IM and EC cell clusters. **i)** Gene expression along IM and EC cell clusters in a) as well as endothelial cells. **j)** Ligand-receptor interactions predicted between endothelial cells and cells of the osteogenic lineage.

Within the two newly defined trajectories for EC and IM osteogenesis, we determined dynamic expression and modules associated with pseudotime as they converge (Fig. 3a-b). Common TFs (*RUNX2, DLX5, SP7, SATB2)* mediating osteogenesis were upregulated toward terminal states across both pathways. The newly described SutureMes1/2 was enriched for osteogenic genes (*SPARC, COL1A1, GAS1)* early in the trajectory and signatures associated with suture formation and regulation in the fetal and postnatal mouse suture (*PRDM6, MSX1)* were most regulated upon transition to SutureMes2 ^55^. Inhibitory control of osteogenesis differed across the pathways, *LMX1B* a common repressor, was enriched late in progenitors across both pathways whereas *TWIST1* was only enriched late in IM progenitors, signifying additional inhibitory regulation in the IM pathway. *NFATC2*, a chondro- and osteo- genesis repressor ^56,57^, was observed early in the IM pathway, but upregulated late in the EC pathway, potentially reflecting context and timing dependent roles in maintenance of the suture niche (IM), and cartilage primordia replacement (EC), respectively. In IM progenitors, accessibility across activators (*RUNX2*) and repressors (*TWIST1, LMX1B*) was simultaneously high at early parts of the trajectory, suggesting they are poised for osteogenesis but remain repressed in the early stages (Fig. 3c). In contrast, at the transcriptome level, there was a reciprocal relationship between repressor and *RUNX2* target expression across pseudotime, suggesting additional, redundant layers of regulation limiting RUNX2 target expression. In the EC pseudo-trajectory, repressor (*TWIST1, NFATC2*) accessibility peaked toward Perichondrium and then decreased, whereas *RUNX2* target accessibility increased as Perichondrium formed, suggesting the latter was critical in driving progression. This contrasts with the gradual reduction in IM repressor accessibility marking the transition into *HHIP+*PreOB alongside a persistently accessible activator (*RUNX2)* (Fig. 3c). In concert, the multi-omic dynamic changes in the IM pathway, in contrast to EC osteogenesis, suggest additional layers of non-transcriptional regulation.

Following suture formation, progressive waves of oriented differentiation emanate from the cranial sutures toward the developing bone front ^45^. In order to identify the newly defined cell states in this progression (stage 2), we utilise OrganAxis (see Methods) to define a maturation axis spanning the coronal suture into regions of the maturing frontal bone. Subsequently, we create zonal bins within this continuous axis based on histological features, and evaluate cell-state mapping along the anterior-posterior (AP) axis (Fig. 3d). Enrichment of *TWIST1*^+^ SutureMes1/2 were substantial in the suture zones (1-3) which constitute the centre of the coronal suture (Fig. 3e). Their enrichment peaked within the osteogenic front where histological features of osteoprogenitors began to emerge along with *HHIP*^+^ PreOB. Establishment of the osteogenic zones coincided with downregulation of anti-osteogenic (*LMX1B*, *TWIST1*), and upregulation of pro-osteogenic (*RUNX2*, *DLX5*, *SP7*) TFs (Fig. 2i-j, 3d-e), signifying a spatially restricted molecular switch that defines territories of the suture. Osteoblast-specific genes including *IRX5*, *SOST*, *SPP1*, *MMP9* and *DMP1* gradually peaked toward the most distant osteogenic zone^58^. Clustering of the spatial voxels aligned osteogenic TFs with axis values away from the coronal suture (Extended Data Fig. 5c).

In EC ossification, bone formation initiates in the interfaces of the synovial joint, with perichondrium formation surrounding the hypertrophic chondrocyte template, giving rise to osteoblast in fetal stages and in postnatal injury repair ^11,52,59^. At the border of the sphenoid in the anterior skull base, we identified the anterior boundary between EC and IM osteogenesis at the frontal-sphenoid interface, with co-localisation of SutureMes1 and SutureMes2, which precede *HHIP*+PreOB and Osteoblast (Fig. 3e), forming a previously undescribed boundary-defining niche. In the limb, we applied ISS-patcher to the ISS data, and inferred spatial trajectories for differentiation of the LimbMes towards perichondrium and osteoblasts in a centripetal pattern in the incipient bone at ∼7 PCW (Fig. 3f). In the older 9 PCW skull base, a comparable pattern was detected in the sphenoid where hypertrophic chondrocytes were surrounded by perichondrium (Fig. 3g).

To further resolve the endothelial-bone relationship within the IM niche, which remains poorly understood compared to the canonical vascularisation models described in the EC niches^60^, we leveraged both the spatial locations and inferred osteo-endothelial interactions to elucidate a plausible cellular basis for endothelial sprouting. As expected within the IM niche, *MSR1*^+^ Tip cells, Mural cells and Capillary Endothelial cells (ECs) progressively co-enriched along the osteogenic zones with Osteoblast and Osteocytes (Fig. 3e, Extended Data Fig. 5a-b), suggesting a spatial-temporal association between vascularisation and osteogenesis. Concurrent to this trend, *ATF4*, a regulatory gene that promotes bone angiogenesis in development ^61^, was also enriched along the axis.

Pro-angiogenic factors *VEGFA* and *VEGFB*, the concentration of which modulates vessel sprouting, were highly enriched at the osteogenic front, which co-localised with SutureMes1/2 and *HHIP*^+^PreOB, which highly expressed *VEGFA* and *VEGFB*, suggesting they are the likely source. Endothelial populations highly express the *VEGF* receptor genes (*FLT1*, *KDR*, *NPR1*) (Fig. 3h), which therefore suggests the former plays a pivotal role in promoting vascular invasion into the bone in IM akin to the well-described function of hypertrophic chondrocytes in EC niches^62,63^.

To determine distinct cell-cell interactions across the EC and IM niches, we utilised NicheNet to compare differences in signalling during sprouting angiogenesis, focusing on recipient tip cells that lead the vascular sprouts. SutureMes1/2 distinctly enriched for *EPHB2* and *EPHB3*, which promotes sprouting behaviour, motility, and vessel formation (Fig. 3i) ^64^. *RACK1*, an intracellular scaffold protein that promotes *VEGF*-*FLT1*-dependent cell migration^65^, was exclusively expressed in tip cells in the calvaria, signifying a highly motile state. *CXCR4*, which promotes sprout anastomosis, was upregulated in tip cells^66^. In EC ossification, tip cells distinctly express *CDON*, the receptor for *IHH* which promotes endothelial proliferation, migration, and angiogenesis. Together, these data support the essential role of SutureMes1/2 and *HHIP*^+^PreOB in coordinating blood vessel invasion and migration in the IM osteogenic niche.

Next, we sought to unravel inferred signalling interactions from the invading endothelial cells toward osteogenic populations. We first defined a spatial gradient of angiogenesis along the defined bone-maturation axis by scoring for enrichment in sprouting angiogenesis pathways (Fig. 3e), which corroborated the coupling of angiogenesis and osteogenesis, orchestrated in the direction of bone growth. We then used CellphoneDB to predict significant signalling interactions from ligands in the endothelial populations toward the recipient osteogenic cell states (Fig. 3j)^67^. Endothelial cells, including capillary ECs, mural cells and tip cells were predicted to signal to osteoblastic cells via DLL1/DLL4–NOTCH2 and JAG1/JAG2-NOTCH2 interacting pairs, which have been reported to promote the differentiation of postnatal perivascular osteoprogenitors^68^. Of note, no *NOG* expression was observed in endothelial cells, unlike the previously described postnatal bone-associated type H vessels^68^, which suggests a novel and unique pattern of crosstalk between endothelial and osteoblastic cells via NOTCH signalling in the embryo. Additionally, mural and capillary ECs promoted osteoblast differentiation through previously described interactions including FGF2-FGFR2^69^ and THBS1-CD36^70^. Mural cells are also likely to be promoting bone mineralization via activation of canonical WNT signalling through the observed RSPO3-LGR5 pairs^71^. In addition to pro-osteogenic roles, the endothelial cells all support osteoclast recruitment and differentiation via CCL14-CCR1^72^ and CXCL12-DDP4 interactions^73^. Through these spatially defined interactions, we discern a model of the interplay between angiogenesis and osteogenesis in the intramembranous niche of the frontal bone whereby tip cells are first recruited by VEGF and EPHB2 from suture mesenchyme and pre-osteoblastic cells, toward the pre-osteogenic condensation that form the bone-front. Vascularising endothelial cells then promote osteoblastic differentiation, osteocyte mineralization and osteoclast recruitment in the maturing bone (Extended Data Fig. 5e).

Other lineages, including neurons have also been reported to play a role in modulating osteogenesis^74,75^. Neuronal development and axon guidance modules were enriched in early stages of both osteogenic pathways (Supplementary Table 4), suggesting involvement of chemical cues from axons in niche formation. Specific IM modules were associated with epithelial mesenchymal transition (EMT), hypoxia response, and TGF signalling, whereas EC-enriched modules were related to neuroactive ligand-receptor interactions, axon guidance and development (GSEA, MSigDB Hallmark 2020 gene sets). To understand the potential role of axon guidance in skull formation, we compute gene-scores for axon guidance modules along the spatial trajectories of osteoblastogenesis in the cranium and revealed enrichment in the suture mesenchyme (Extended Data Fig. 5f), consistent with previous description of axonogenesis regulating suture formation in the mouse^76^. Future studies capturing transcriptional profiles of neuronal cell bodies of the innervating axons may shed light on the cell-cell interactions at play.

### Inferring *PAX7*^+^ chondrocyte origins

Chondrogenesis is an essential process for the formation of various types of cartilage, including hyaline, fibrous, and elastic cartilage. To reveal the transcriptional heterogeneity of chondrocytes, the chondrocyte subcompartment was further sub-clustered across anatomical regions and timepoints and annotated based on canonical marker genes (Fig. 4a, b, Extended Data Fig. 6a and Supplementary Table 2). To dissect cellular heterogeneity within a spatial context, we applied ISS-Patcher and Cell2location to transfer cell state labels from our single-cell atlas to ISS and 10x Visium slides from the shoulder at 7.3 PCW and calvaria at 9 PCW (Extended Data Figs. 6b, c).

**Figure. 4:**
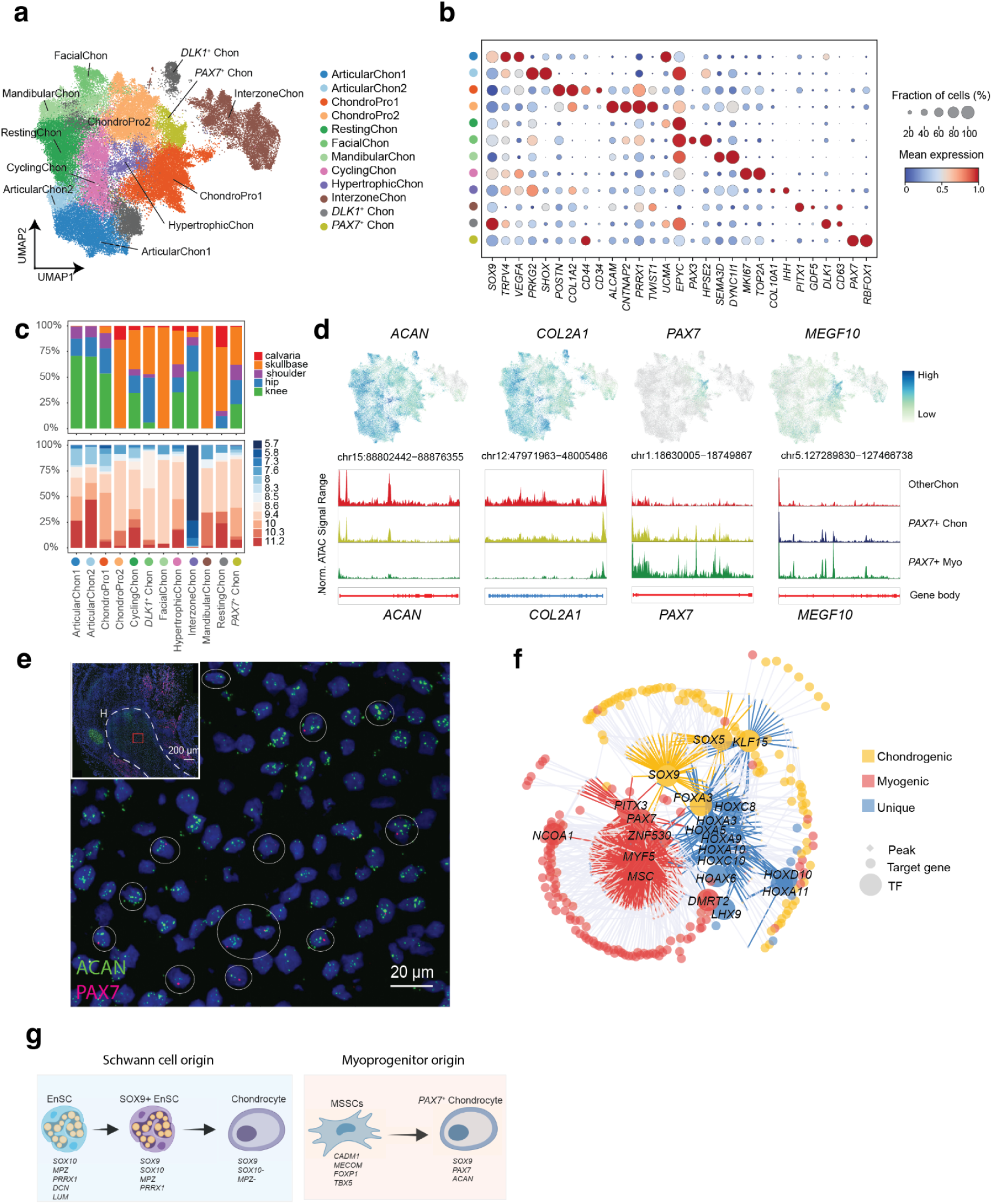
Chondrogenesis across different anatomical regions. **a**) RNA UMAP embeddings of chondrocytes coloured by subtypes. **b**) Dot plot showing the mean expression (dot colour) and fraction of expressing cells (dot size) of selected marker genes. **c**) Distribution of different chondrogenic sub types across anatomical regions. a, b, c share the same legends. **d**) RNA UMAPs (the upper panel) of representative chondrogenic genes (*ACAN*, *COL2A1*) and myogenic (*PAX7*, *MEGF10*) for *PAX7*^+^ chondrocytes. Genome browser tracks (the lower panel) from scATAC-seq analyses displaying the sum signal within the indicated gene loci. OtherChon represents the whole chondro-lineages except *PAX7*^+^ Chon. **e**) RNA scope for cells that co-expressed myogenic markers (*PAX7*, red) and chondrogenic markers (ACAN, green) in the shoulder at 7.3PCW. The blue marks the cell nucleus. Scale bar, 20/200 μm. **f**) Enhancer-GRN showing enriched TFs in *PAX7*^+^ Chon, inferred using SCENIC+. Circles and diamonds represent genes and regions with TF-binding sites, respectively. Colour indicates whether the TF of *PAX7*^+^ Chon is also differentially expressed in chondro-lineages or *PAX7*^+^ myocytes. TF-region links are coloured by TF. **g)** Schematic representation of the putative non-classical lineages that contribute to chondrogenesis.

Our chondrocyte clusters exhibited strong regional-specificity due to divergent development origins (Fig. 4c)^77^. In terms of cell clusters, we identified known populations, such as hypertrophic chondrocyte (HyperChon, *COL10A1*, *IHH*), cycling chondrocyte (CyclingChon, *MKI67*, *TOP2A*), resting chondrocyte (RestingChon, *UCMA*), interzone chondrocyte (InterzoneChon, *GDF5*, *PITX1*), as well as new populations, including two chondrocyte progenitors (ChondroPro1, ChondroPro2), anatomical region-specific chondrocytes, *DLK1*^+^ chondrocytes (*DLK1*^+^ Chon, *DLK1*, *CD63*), and *PAX7*^+^ chondrocytes (*PAX7*^+^ Chon, *PAX7*, *RBFOX1*). Two region-specific ChondroPros, which were enriched in appendicular joints and skull respectively, transitionally expressed fibroblast differentiation markers (*POSTN*, *COL1A1*, *PRRX1*, and *TWIST1*), which was consistent with findings in early chondrocyte progenitors in mice^19^. In the skull base, facial chondrocytes highly express *PAX3*, indicating an origin from neural crest^78^. Mandibular chondrocytes highly expressed *SEMA3D* in posterior regions^79,80^. In appendicular regions, we observed two subtypes of articular chondrocytes, with one population being enriched for *TRPV4* and *VEGFA*, while the other one was more mature, with relatively low *SOX9* expression and high *EPYC* expression. In particular, *DLK1*^+^ Chon was highly enriched in ribosomal genes and distinctly expressed *CD63* and *DLK1*. *CD63* has been identified in the pre-hypertrophic layer in the limb, while *DLK1* is a novel marker for embryonic chondroprogenitor cells undergoing lineage progression from proliferation to pre-hypertrophic stages. Spatially, CyclingChon, *DLK1*^+^ Chon and HyperChon were also arranged sequentially within the bone, spanning from the epiphysis toward to diaphysis, the primary ossification centre (Extended Data Fig. 6b), which corroborates the transitional role of *DLK1*^+^ Chon, from proliferative to pre-hypertrophic phenotypes during chondrogenesis.

With regard to gene regulatory networks, we identified common and region-specific co-expressed gene modules using Hotspot^81^ and performed gene ontology enrichment on each module (Extended Data Figs. 6 d,e). Shared gene modules (M3, M5, etc) across regions mainly related to chondrocyte cellular structure and functional development, including biomineralization, collagen-containing extracellular matrix, and cartilage development. Interestingly, synapse related pathways were enriched both in appendicular and cranial samples, suggesting axons may contribute to development of cartilage^82^ which in pathology may be responsible for joint pain. In addition, chondrocytes from the skull base enriched for axon guidance gene modules, while chondrocytes from appendicular joints had specific gene modules, including glucose metabolism and cellular response to hypoxia (Extended Data Figs. 6f, g). Taken together, our data provides a comprehensive catalogue of chondrocyte molecular profiles in multiple developing human joints.

Moreover, our work uncovered a previously undescribed developmental chondrocyte cluster (*PAX7*^+^ Chon), which first appeared at 7 PCW and peaked between 9-10 PCW across multiple donors (Fig. 4c). *PAX7*^+^ Chon co-expressed markers and gene modules of chondrocytes and muscle cells on the transcriptomic and epigenetic level (Fig. 4d), and showed overlap of genes that enrich in cartilage and muscle development pathways (Extended Data Fig. 6h, i). To interrogate the possibility of doublet-capture artefact, we first applied stringent doublet processing by performing extra statistical tests over the doublet score (see Methods, Extended Data Fig. 7a, b). Then we verified the presence of *PAX7*^+^ Chon through in situ hybridization experiments (Fig. 4e), which was consistent with the multiplexed ISS data (Extended Data Fig. 6b). Additionally, *PAX7*^+^ Chon displayed transcriptional characteristics not explained by doublet formation (Extended Data Fig. 7b,c). To reveal GRNs that govern the transcriptional identity of *PAX7*^+^ Chon, we applied SCENIC+ and constructed a GRN based on differentially expressed transcription factors (TFs) with their predicted target genes. Aside from identifying key myogenic and chondrogenic key regulators (*PAX7*, *MYF5*, *SOX5*, *SOX9*), we also discovered core posterior axis HOX gene modules, suggesting that patterning of major synovial joint chondrocytes may be conferred through their nested HOX expression ^83^.

Previous mouse studies suggested that chondrocytes may share origins with the muscle lineage^84,85,86^. Skeletal muscle mesenchymal cells (SkM.Mesen)^15^ not only express classical muscle markers at low levels, but also exhibit gene expression related to mesenchyme. Myo-skeletal stem-like cells (MSSCs) expressed *CADM1*, *MECOM*, and *FOXP1*, previously reported as makers of mouse skeletal/stem progenitors^87^, along with genes related to muscle, cartilage, and mesenchyme. In addition, MSSCs expressed low levels of lateral plate mesoderm markers. Conversely, *PAX7*^+^ MyoProgenitor cells highly expressed muscle-related genes, suggesting they are a distinct branch separate from *PAX7*^+^ Chon (Extended Data Fig. 7c). We further interrogated the relationship between these progenitors and *PAX7*^+^ Chon, applying scFate, to reveal a trajectory from MSSCs to *PAX7*^+^ Chon, with upregulation of chondrocyte genes and downregulation of stem cell genes, such as *SOX4*, *CDH2* and *CDH11* on both the RNA and ATAC level (Extended Data Figs. 7d-f). Comparison of mouse and human cell clusters revealed similarity between *PAX7*^+^ Chon and a *MYF5*^+^ progenitor-derived mouse cluster from the extraocular muscle (Extended Data Fig. 7g), which was reported to derive from progenitors that give rise to both myogenic and connective tissues in mouse^15,86^. Taken together these results indicate the possibility of a new source of chondrocytes from muscle progenitors, which suggested the potential complexity of embryonic cell types prior to reaching terminal differentiation.

Lineage tracing experiments in zebrafish and mice^88^ have shown that Schwann cells possess the evolutionarily conserved potential to differentiate into chondrocytes during embryonic development in both species. Here, we identified *SOX9*^+^ endoneurial Schwann cells (*SOX9*^+^ enSC) within the Schwann cell compartment, characterised by the expression of typical chondrocyte (*SOX9*, *COL9A1*, *ACAN*, *COL2A1*), and classical Schwann cell markers (*MPZ*, *SOX10*) (Extended Data Figs. 8a-c). Velocity analysis suggested that multipotent hub Schwann cells (Hub SCP) serve as the foundational cells in the differentiation hierarchy. *SOX9*^+^ enSC represented one of the endpoints on the mesenchymal trajectory, expressed a mesenchymal gene signature (*PRRX1*, *PRRX2*, *PDGFRA*, *TWIST2*) and upregulated *HOX* TFs (*HOXA9*, *HOXA10*, *HOXA11*, *HOXD10*) (Extended Data Figs. 8d, e). We applied RNAscope (*SOX9*, *SOX10*, *MPZ*) to localise *SOX9*^+^ EnSC in the acetabulum at 7.3 PCW (Extended Data Figs. 8g, h). In summary, the analyses suggest that Schwann lineage cells may carry the potential to serve as an alternative source of chondrocytes in humans. However, further lineage tracing experiments are essential to validate this. We therefore hypothesise that chondrocytes derive from multiple potential origins, such as muscle and Schwann lineages during embryonic development (Fig. 4i).

### Developmental links to complex traits of the adult skeleton

Numerous conditions of the postnatal and ageing skeleton have been linked to disrupted joint and bone changes during the embryonic stages of life. Of note, enhancer-associated variants linked to adult osteoarthritis (OA) appear to act on anatomical-region-specific regulatory networks to influence synovial joint morphology during development^89^. However, the cell states that mediate this process remain evasive.

To explore potential causal links in foetal skeletal cell states for adult complex traits, we first conducted fGWAS (Fig. 1c, see Methods), which integrates full GWAS summary statistics with cell-state specific epigenetic and transcriptomic signatures across clusters, computing cell-cluster enrichments for phenotypes. Interestingly, we found distinct lineage enrichments for knee and hip OA in chondrogenic and osteogenic cell populations, respectively (Fig. 5a). Knee OA and its surrogate phenotype, total knee replacement (TKR) enriched in most chondrogenic states, except for *PAX7*^+^ chondrocytes and InterzoneChon (*GDF5*^+^). In contrast, hip OA enrichment was only significantly observed in two chondrocyte populations (ChondroPro and Hypertrophic Chondro), but was additionally enriched in Perichondrium, Osteoblast as well as Tenocyte. We further performed fGWAS on our fine-grained annotations (Fig. 5b) to explore subcluster enrichment. Consistent with the broad annotations, various types of chondrocytes show significant enrichments for knee OA (Fig. 5b) whereas the signal for InterzoneChon implicated Early IZ in hip OA. These findings implicate a greater impact for bone development affecting hip OA risk, potentially through modulating hip shape and a thereby altered risk of mechanical stress at the hip joint over time^90^. In contrast, the knee-specific findings point toward the central role of chondrogenesis in knee pathology, potentially implicating postnatal regenerative mechanisms.

**Figure 5:**
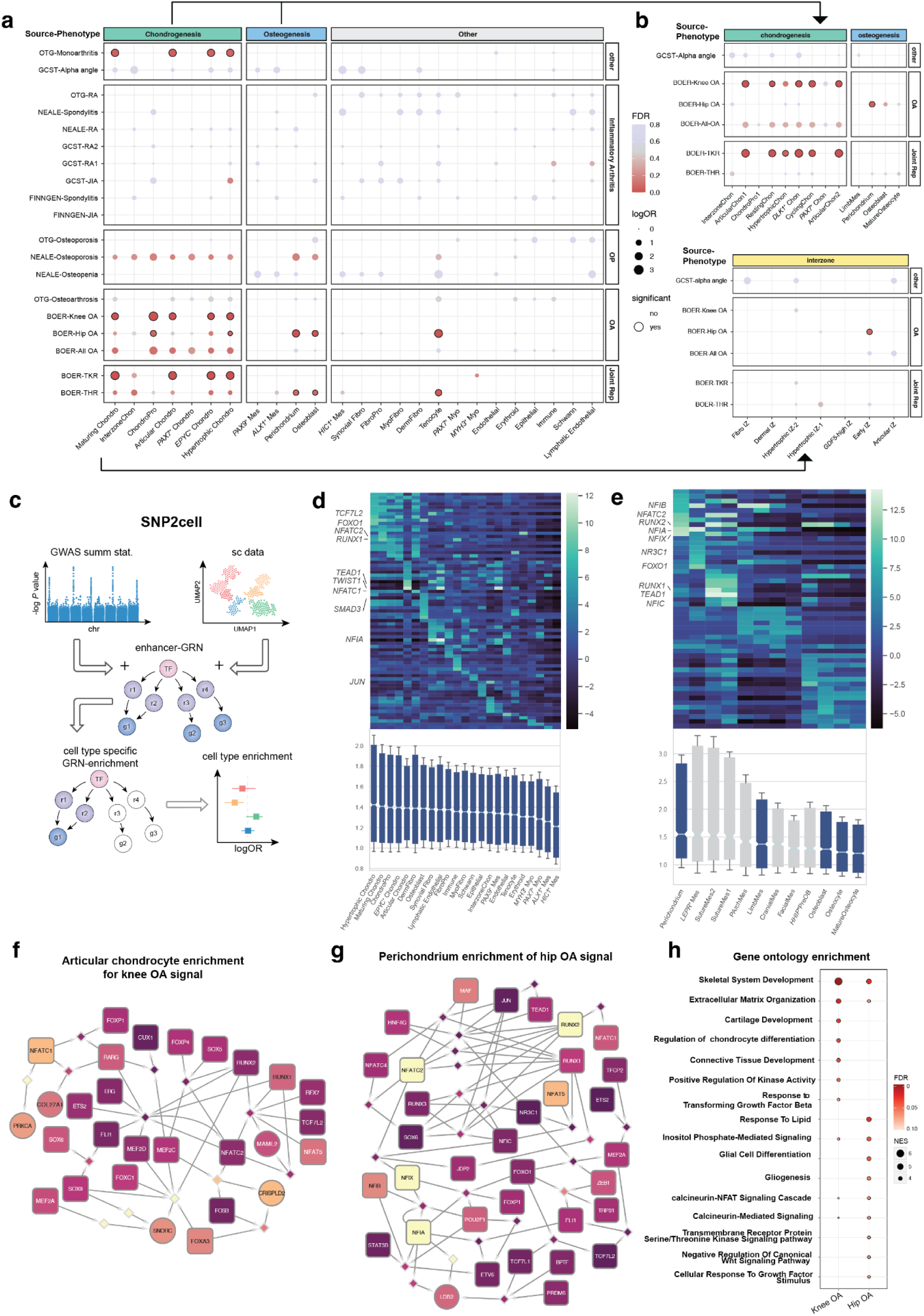
Links to complex diseases of the skeleton. **a)** fGWAS analysis results for enrichment of GWAS signals for bone-related complex traits in broadly annotated cell states. **b)** Enrichment of selected complex traits in finely annotated osteo- and chondrogenic cell states (top) and InterzoneChon subclusters (bottom). **c)** Schematic representation of SNP2cell: scores derived from GWAS summary statistics and cell cluster marker scores are mapped and integrated across a gene regulatory network, highlighting enriched modules predicted to play a cluster-specific role in disease. **d)** Enrichment scores for knee OA across all cell clusters (heatmap, top) showing a cluster of similar enriched genes for chondrocytes and median enrichment scores (boxplot, bottom. center line, median; boxes, first and third quartiles of the distribution; whiskers, highest and lowest data points within 1.5 × interquartile ratio (IQR)). **e)** Enrichment scores for hip OA across the osteogenic trajectory (heatmap, top) and median enrichment scores (boxplot, bottom . center line, median; boxes, first and third quartiles of the distribution; whiskers, highest and lowest data points within 1.5 × IQR). **f)** Articular chondrocyte and knee OA specific enriched sub-network. Brighter colours correspond to a larger enrichment score, relative to scores obtained from random permutations. **g)** Perichondrium and hip OA specific enriched sub-network. Brighter colours correspond to larger enrichment scores as in d. **h)** Gene Set Enrichment Analysis for GO Biological Process terms across Articular Chondro-knee OA and Perichondrium-hip OA enrichment scores.

Complex diseases are often associated with many mutations across non-coding regions. We therefore sought after a way to integrate the GWAS signals of hip and knee OA described above across enhancers of our gene regulatory network to identify enriched TFs and pathways, while accounting for cell cluster specific effects. To this end we developed SNP2CELL, a tool that uses gene regulatory networks as a basis to aggregate scores of individual SNPs and cluster marker genes across pathways of connected TFs (Fig. 5c). Aggregation of scores was achieved through a network propagation approach^91^. In contrast to simply linking SNPs with their closest genes, this also allows for the identification of upstream TFs which might not be mutated, but whose effect on multiple binding sites and target genes is often disrupted, by integrating a large number of disease associations. By using eGRNs previously computed with SCENIC+, we can utilise SNP signals mapped to both gene regions and distal enhancers in ATAC peaks. By overlapping integrated SNP scores with integrated marker gene scores per cell cluster, and comparing the effects to random permutations, we obtain cell cluster specific enrichments of sub-networks of our initial eGRN.

Deriving insights from cell states implicated in fGWAS, we next clustered enrichment scores for knee and hip OA, focusing on osteogenesis for the latter. In knee OA, various chondrocyte subtypes clustered together by high enrichment score and average gene enrichments (Fig. 5d), mirroring its fGWAS enrichment for chondrocytes. Likewise for hip OA, Perichondrium expectedly had the largest average enrichment across the osteogenic trajectory (Fig. 5e) consistent with fGWAS results. Utilising SNP2CELL, we identified sub-networks for knee OA and hip OA, prioritising articular chondrocytes (Fig. 5f) and perichondrium (Fig. 5g), respectively, and revealing similar regulatory pathways that balance chondro and osteogenic functions.

In the knee articular chondrocyte knee OA network (Fig. 5f), a number of non-TF genes with roles in cartilage makeup and chondrocyte differentiation (*COL27A1*, *PRKCA*, *SNORC*, *CRISPLD2*) were identified and predicted to be regulated by *NFATC1* and *FOXA3*. *COL27A1*, and the proteoglycan *SNORC*, are involved in chondrocyte ECM makeup and maturation^92^. However, *PRKCA* is a kinase that has been linked to mechano-sensing in articular chondrocytes^93^. Notably, *NFATC1* itself has been described as a marker of articular cartilage progenitors affecting differentiation^94^ and also protects against OA^95^. For hip perichondrium in hip OA (Fig. 5g), the osteogenic key regulator *RUNX2* showed significant enrichment, along with multiple *NFAT* genes (*NFATC1*, *NFATC2*, *NFATC4*, *NFAT5*), which in conjunction with additional TFs (*ZEB1*, *MAF*, *TEAD1*) implicated calcineurin and WNT signalling pathways. Both canonical and non-canonical WNT signalling via Ca^2+^/Calcineurin/NFAT have been linked to bone formation and remodelling, affecting the balance and differentiation of both osteoblasts and osteoclasts ^96^. Various WNT signalling inhibitors including *DKK1* and *FRZB* have also been linked to the shaping process of the hip and osteoarthritis, which may be particularly important during foetal development^97^. To provide a summary of the potential functional role of the enriched disease-associated cell-cluster specific genes, we conducted gene set enrichment analysis (GSEA) on Gene Ontology (GO) terms. This expectedly featured terms related to extracellular matrix organisation, cartilage development and chondrocyte differentiation in articular chondrocytes in knee OA specifically, whereas hip perichondrium scores showed enrichment for inositol-phosphate, calcineurin and NFAT signalling, and cellular response to lipid (Fig. 5h). The latter points to the potential interplay of lipids with bone formation and osteoarthritis^98^. Overall, through application of fGWAS and our newly developed pipeline SNP2Cell, we were able to distil differential involvement of the chondrogenic and osteogenic lineages respectively in hip and knee OA, and provide a cellular basis for the link between developmental cell states and regionally distinct adult disease.

### Deciphering monogenic conditions of the embryonic skeleton

Both syndromic and non-syndromic forms of craniosynostosis are congenital conditions that involve disturbances in cranial ossification and suture formation during fetal and postnatal development. Disease-associated clinical features include premature cranial suture fusion, and depletion of osteoprogenitor pools that facilitate the postnatal expansion of the cranium, leading to severe global developmental consequences. Disease mechanisms that underlie these changes are reportedly linked to missense mutations and haploinsufficiency in genes that govern persistence of IM osteo-progenitor pools within the suture joints throughout the cranium ^99–101^.

To implicate cell states that underlie craniosynostosis disease mechanisms we first cross-referenced pseudotime-associated DEGs enriched in the IM and EC pathways against a candidate database of over 2700 genes known to cause congenital conditions in humans^11^. We then filtered for enriched DEGs that were associated with musculoskeletal (MSK) conditions and created a global view of enriched expression and accessibility across the intersecting genes (Fig. 6a). Within these DEGs, we then filtered for craniosynostosis associated genes by combining a subset of the database, and reported genes obtained from Genomics England Limited rare and common craniosynostosis panels (Supplementary Table 5), of which a high proportion of the top DEGs were associated with synostosis (Fig. 6a). Most of these associations (n=13/22) were within the progenitor populations of the IM trajectory (SutureMes1/2 and *HHIP*+PreOB). Broadly, most of the craniosynostosis enriched genes were highly accessible and expressed in IM progenitors, apart from *FN1* and *IHH*, suggesting the embryonic period is critically affected in craniosynostosis.

**Figure 6:**
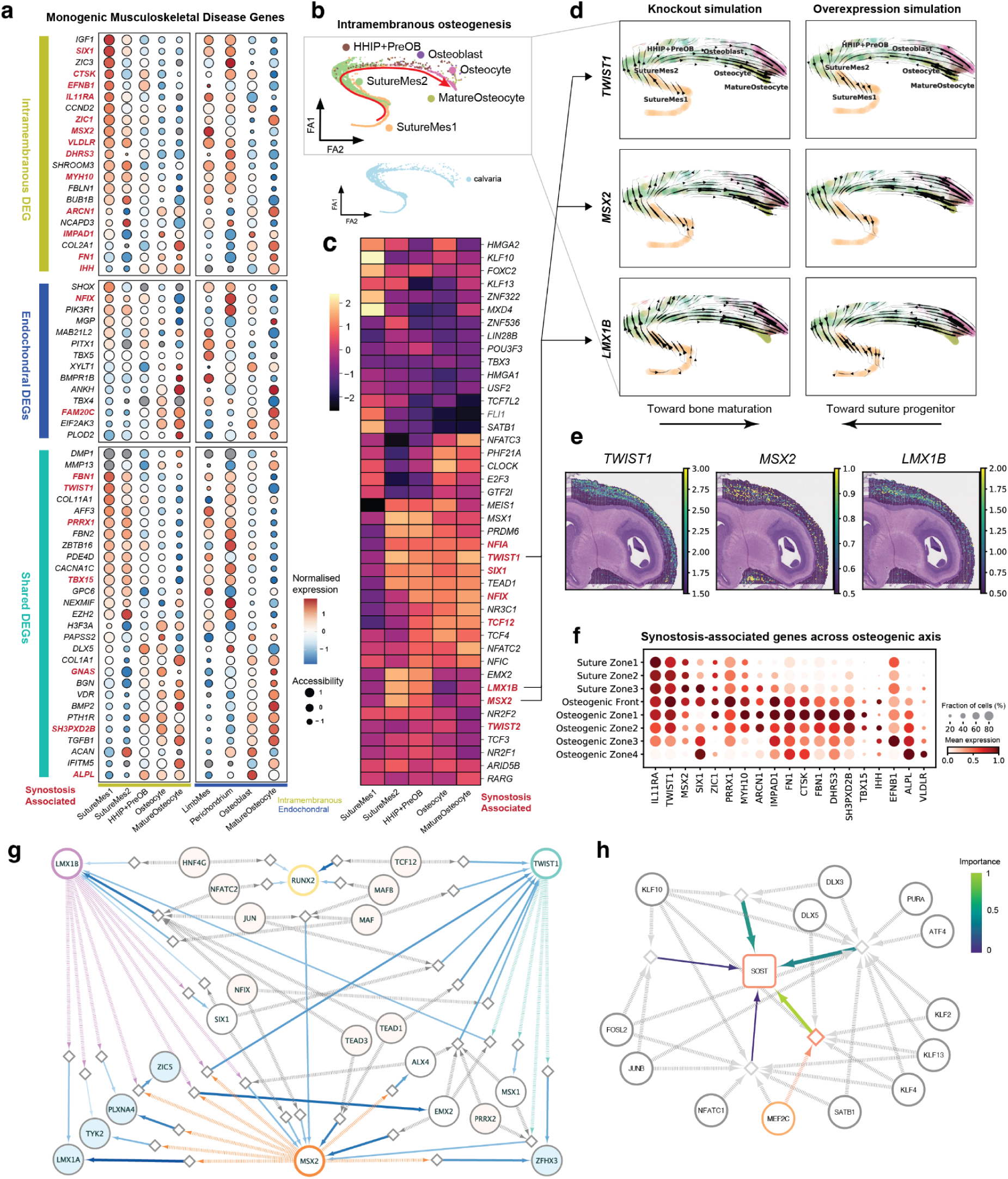
Links to Monogenic conditions affecting bone development. **a)** Known genes mutated in musculoskeletal monogenic conditions with their expression and accessibility along the intramembranous and endochondral osteogenesis trajectory. **b)** Force-directed embedding of the endochondral trajectory showing cell cluster annotations. **c)** Heatmap of *in silico* TF knockout perturbation scores per cell cluster. A higher score indicates transcriptomic changes induced by the perturbation are promoting osteogenesis. **d)** TF perturbation vectors showing the direction of induced transcriptomic changes on the FA embedding for three selected TFs: TWIST1, MSX2 and LMX1B. KO simulation of all three genes promotes osteogenesis, while overexpression inhibits it. **e)** TF expression on Visium Cytassist of the frontal region of skull, enrichment of TFs in coronal suture and skull-forming layers. **f)** Expression of synostosis-associated genes along spatial zones defined using OrganAxis. **g)** Enhancer-GRN showing inter-regulation between TWIST1, MSX2 and LMX1B, inferred using SCENIC+. Circles and diamonds represent genes and regions with TF-binding sites, respectively. Region-gene links are coloured and scaled according to peak2gene importance, while TF-region links are coloured by TF. Blue circles represent regulated targets, orange circles regulators and white intermediate genes. **h)** Enhancer-GRN showing predicted regulation of SOST. Region-gene links are coloured and scaled according to peak2gene importance. Highlighted in red is a region containing the enhancer known to be mutated in van Buchem disease.

To elucidate potential effects of the synostosis-associated TFs on gene changes during osteogenesis, we applied CellOracle to predict velocity shifts within the newly defined trajectory and eGRNs using *in silico* perturbation simulation of gene knockouts in the IM osteogenesis sub-compartment (Fig. 6b-c). Of the top-genes predicted to cause greatest velocity shifts in the trajectory, *TWIST1*, *MSX2*, *LMX1B* were all disease-associated genes that lead to highest velocity shifts in SutureMes2 when perturbed (Fig. 6c). The former two were also IM pseudotime-associated DEGs (Fig. 6a), suggesting the velocity shift is highly likely to reflect the loss-of-function mechanism of disease pathogenesis in these TFs ^102,103^. Visualisation of the inferred velocity shifts revealed expected patterns of knockout favouring lineage progression and bone formation (Fig. 6d), which are likely to penetrate in regions of the coronal suture that indeed correspond to spatial enrichment of the candidate TFs (Fig. 6e). Numerous other disease-associated TFs demonstrated maximal enrichment in the Suture Zones (*IL11RA*, *SIX1*), whereas others preferentially affected more mature parts of the trajectory (*IHH*, *ALPL*, *VLDLR*) (Fig. 6f). Overall, these findings constitute novel explanations for features of TF-mediated craniosynostosis including premature suture fusion and ossification.

To explore potential cell-extrinsic influences on fetal bone development, we curated a list of 65 clinically approved drugs from the chEMBL database which carried warnings of teratogenicity (Supplementary Table. 6). We then utilised to score target gene-group expression per cell state within our osteogenic sub compartment. Interestingly, this revealed overall greater target enrichment of teratogenic drugs within IM progenitors and downstream osteoblast/osteocyte cell states compared to EC progenitors (Extended Data Fig. 9a). Retinoid drugs expectedly enriched for targets in Osteocytes, consistent with observations of retinoic acid enhancing *in vitro* osteogenesis in mouse suture cells^104^. Likewise, the well-known skeletal development teratogen thalidomide enriched for targets in Osteocytes. Aside from antihypertensives targeting the renin-angiotensin pathway (*ACEi*, *ARB*, Renin inhibitor) which enriched for targets in SutureMes1 and *HHIP*^+^PreOB, we also identified high enrichment for endothelin receptor antagonists (ERAs) specifically in SutureMes1. Mice lacking the endothelin A receptor recapitulate wide-spread cranial neural crest-related defects in utero^105^. Informed by teratogenic effects of ERA use in animals^106^, they are not routinely prescribed for pulmonary arterial hypertension (PAH) during pregnancy. However, there remains a lack of evidence for teratogenicity in humans, and women of child-bearing age continue to utilise them for treatment of PAH. Our analyses support the notion of potential targets for ERAs within the human suture cells and lend support to a contraindication for their use during pregnancy.

To further resolve the regulatory relationship between the prioritised TFs (*TWIST1*, *MSX2*, *LMX1B*) within the osteo-lineage, we utilised our paired snRNA-seq and scATAC-seq dataset to reconstruct interaction networks across the TFs (Fig. 6g). Our predictions revealed a rich web of inter-regulation of shared coding and non-coding targets across the three candidate TFs. Of these connected nodes, numerous TFs are associated with loss-of-function mutations leading to craniosynostosis (*SIX1*, *TCF12*, *NFIX*, *ALX4*), suggesting a tightly regulated network conferring suture patency, inhibiting osteogenesis in this region. Notably, *TCF12* was reported to govern normal coronal suture development through heterodimer formation with *TWIST1*, and severe phenotypes are observed in mice with doubly deleterious mutations^102^. *TCF12* served as a node connecting *TWIST1* to *RUNX2*, suggesting that disruption of this critical co-regulatory network leads to human disease. While direct transcriptional regulations of *RUNX2* by the three inhibitory TFs were predicted to be weaker than their stronger inter-regulation and not visualised in Fig. 6g, predicted regulations of *RUNX2* (Extended Data Fig. 9b) therefore likely contribute to known complex regulatory mechanisms, including protein-level interactions^107^. In addition, cell-extrinsic signals and control mechanisms are likely to act on the connecting nodal points such as *NFATC2* ^96^, governing the balance of osteogenesis.

Multi-omic information additionally allows exploration of enhancer-mediated regulation of transcription. This is of particular importance for monogenic bone-forming conditions such as Van Buchem disease (VBD) whereby mutation in the non-coding ECR5 enhancer of *SOST* leads to sclerosing dysplasia of bone^108^. We therefore constructed a regulatory network centred around *SOST* and identified the ECR5 enhancer to directly regulate SOST (Fig. 6h). Numerous osteogenesis-associated TFs were predicted to regulate the region containing the ECR5 enhancer (*DLX5*, *KLF2*, *KLF4*, *KLF13*, *MEF2C*). Notably, *MEF2C* has previously been shown to regulate ECR5 in mice^109^, confirming its direct role in controlling *SOST* transcription. Overall, these predictions form a basis for constructing cellular models of TF and enhancer-driven monogenic conditions of the bone lineage.

## Discussion

We present the first multiomic cell atlas to capture the spatially-resolved cellular composition of bone, synovial and suture joint development across the first trimester in humans. First, we dissected the cellular taxonomy of interzone cells, including *GDF5*-expressing progenitors in the synovial joints, and developed ISS-Patcher, a tool to aid inference of IZ subclusters in our high-resolution 155-plex ISS data. We highlighted Articular, Fibro (*GDF5*^+^) and (*GDF5*^-^) Hypertrophic IZ subclusters which zonate the articular chondrocyte and cartilage template-forming boundaries of the embryonic synovial joints, respectively.

Within lineages described in mice, *GDF5*^+^ cells are posited to give rise to the numerous cell components within the joint including ligaments and tendons. Here we describe a novel two-stage process whereby the fibroblasts and tenocytes of the synovial joints arise from embryonic *HIC1*^+^ fibroblasts, a population previously reported in the mouse embryonic limb^29^, and a fetal (*PI16*^+^) fibroblast progenitor which is transcriptionally similar to human postnatal universal fibroblasts ^30^. Together, these shed new light on the cellular architecture of the developing human embryonic synovial joint in the first trimester.

We have identified numerous previously undescribed human cranial embryonic osteo-lineage cell states including CranialMes (*HAND2*)^110,111^, FacialMes (*PAX3*)^51^ and PArchMes (*LHX8*)^112^, by their expression of comparable marker genes in mice counterparts. We leveraged our spatial analyses of the fetal cranium to define two novel fetal *TWIST1*^+^ SutureMes populations, and *HHIP*-expressing pre-osteoblast which contributes to osteogenesis, and mirrors populations of the fetal mouse suture ^30,54^. We also discovered distinct IM and EC osteogenic cellular trajectories in their respective regional niches. Using OrganAxis, we illustrate the spatial component of these trajectories as they develop and show a regulatory shift in the poised osteogenic suture population as it progresses in space toward maturing pre-osteoblastic cell states where we uncover a region that critically recruits endothelial cell states to drive ossification. We show that the EC niche in the appendicular anlagen derives from an undifferentiated progenitor expressing *ISL1* and *TBX5*, which mark an equivalent population in the human limb ^11^, and proceeds to form perichondrium and osteoblasts through developmental time and space in the first trimester. These novel cellular trajectories form a valuable reference for future studies of bone development. Further studies of the second trimester, when adipose cell states and mature ligamentous tissues develop, will facilitate a complete census of the joint.

We lend support to a potential myoprogenitor-related origin for the newly reported *PAX7*^+^ chondrocyte, and likewise suggest that a novel *SOX9*^+^ Schwann cell state may confer chondrogenic potential. These proposed cross-lineage cell fates uncover potential similarities to mouse development described above, and crucially, the value of computational tool development in unravelling comparative human biology. In the future, more focused investigations that involve isolation of these cell lineages will further inform their biology.

Leveraging our dataset, we were able to identify developmental links to both monogenic and complex musculoskeletal diseases. We observed marked differences in the enrichment of hip and knee OA GWAS signals across cell clusters, implicating bone formation in the former, and chondrogenesis in the latter, which we traced back to the osteogenic perichondrium, and numerous chondrogenic cell states, respectively. In future, this framework may be applied to other conditions to gain insights into cell-state specific pathogenesis. Lastly, by systematically simulating in silico KO of TFs known to be associated with craniosynostosis, we identified a network of regulators inhibiting osteogenesis, which provides mechanistic detail of how previously reported monogenic loss-of-function mutations in, for example, *TWIST1*, *MSX2* and *LMX1B*, may lead to early suture fusion. This approach, applied to the newly mapped cross-region osteogenic trajectory, holds immense value in predicting potential effects of gene changes across other diseases involving the cellular lineage.

Our comprehensive multimodal human fetal skeletal atlas is a unique fundamental resource for the understanding of human cartilage and bone development in the first trimester. It also has the potential to serve as a cellular roadmap for *in vitro* endeavours to differentiate osteoblast and chondrocyte and mesenchymal cell states in the developing musculoskeletal system.

## Acknowledgements

We thank the donors and their relatives who contributed to this study. We thank the staff of the Teichmann group wet lab team, Cellular Genetics Informatics team, Cytometry Core Facility team, Core DNA Pipelines team, and Spatial Genomics Platform team at the Wellcome Sanger Institute for their support. We also thank M. Storer, S. Kimber, J.E.G. Lawrence, A. Maartens for assistance in manuscript writing, A. Oszlanczi and A. Wilk for sample management and administrative assistance, C. Battrum for administrative assistance and S. Davidson for helpful discussions.

This work was made possible in part by the Wellcome Trust (220540/Z/20/A, 221052/Z/20/Z, 221052/A/20/Z, 221052/B/20/Z, 221052/C/20/Z, and 221052/E/20/Z); the NIHR Cambridge Biomedical Research Centre (NIHR203312) – (The views expressed are those of the authors and not necessarily those of the NIHR or the Department of Health and Social Care); the MRC Clinical Research Training Fellowship to K.T; the Wellcome Trust Sanger Institute Postdoctoral Fellowship to LF; This project has received funding from the European Union’s Horizon 2020 research and innovation programme under the Marie-Skłodowska-Curie grant agreement No. 101026233 to J.P.P.

## Author contributions

K.T., L.F. AND J.P.P. conceived the study, performed data generation and experiments, conceived methodology, performed analysis, collected biological samples, and wrote the paper. K.R. performed data generation, conceived methodology, collected biological samples, performed analysis, conducted imaging, and wrote the manuscript. T.L., N.Y. performed computational analysis and conceived methodology. P.H. performed computational analysis and provided key biological interpretation. C.X. performed computational analysis and provided key biological interpretation. J.C. performed experiments, conceived methodology, and collected samples. R.L.. performed computational analysis and provided key biological interpretation. K.K. performed computational analysis and provided key biological interpretation. N.H. performed analysis and provided key biological interpretation. S.M. performed analysis and provided key biological interpretation. L.R., R.K., S.P., A.W.C. performed experiments and sample processing and collection, generating single-cell data. E.T, I.M. and F.M. performed imaging experiments, provided key biological insights, and collected tissue samples, and generated spatial data. B.C., A.V.P., D.H., S.M., M.P. and P.M. performed computational processing of data and performed data analysis. X.H. conducted tissue processing and sample collection. K.B.M. conceived computational methodology and provided key biological interpretations. M.H. conceived the study and provided key biological interpretations. R.A.B. conceived the study and provided key biological interpretations. O.B. conceived the imaging experiments, provided key biological interpretations and analysed the data. C.D.B. conceived the study, conceived methodology, provided key biological interpretation, and supervised the project. S.A.T. conceived the study, conceived methodology, provided key biological interpretation, and supervised the project.

## Data availability

High-throughput raw sequencing data in this study is available from ArrayExpress (www.ebi.ac.uk/arrayexpress) with accession numbers. Processed snRNA-seq/scATAC-seq data, visium and ISS data are available for visualisation and downloading from https://developmental.cellatlas.io/skeleton-development

## Code availability

ISS-Patcher is available at https://github.com/Teichlab/iss_patcher. SNP2CELL is available at https://github.com/Teichlab/snp2cell.

## Materials and Methods

### Sample acquisition and ethics

Developing human limb and cranium tissue samples were obtained from elective terminations under REC 96/085 (East of England - Cambridge Central Research Ethics Committee). Briefly, samples were kept suspended in PBS and at -4°C on ice during dissection. Shoulder, hip and knee joints were dissected en-bloc from the limbs. For the shoulder joint a proximal incision was made at the distal third of the clavicle, and a distal incision was created at the neck of the humerus. For embryonic shoulder samples where distinctive bone features had not formed, approximations were made to capture the entirety of the glenohumeral and acromioclavicular joints. For the cranium samples (<8 PCW), two regions were dissected for each of the calvaria and skull base, separated at the posterior border of the frontal bone in both cases. For older cranial samples (>8 PCW), tissues were dissected to separate the frontal, parietal, sphenoid, ethmoid, occipital and temporal bones where feasible. Samples were initially embedded in optimal cutting temperature medium (OCT) and frozen at -80°C on an isopentane-dry ice slurry. Cryosections were then cut at a thickness of 10 μm using a Leica CM1950 cryostat and placed onto SuperFrost Plus slides (VWR) for ISS or Visium CytAssist, or used directly for single-nuclei processing.

### In situ sequencing (ISS) and high-resolution imaging

ISS was performed using the 10X Genomics CARTANA HS Library Preparation Kit (1110-02, following protocol D025) and In Situ Sequencing Kit (3110-02, following protocol D100), which comprise a commercialised version of HybRISS^113^. Probe panel design was based on fold-change thresholds in cell states of the limbs (Supplementary Table 3). Briefly: cryosections of developing limbs were fixed in 3.7% formaldehyde (Merck 252549) in PBS for 30 minutes and washed twice in PBS for 1 minute each prior to permeabilization: sections were briefly digested with 0.5 mg/ml pepsin (Merck P7012) in 0.1 M HCl (Fisher 10325710) at 37°C for 15 seconds (5 PCW) or 30 seconds (6 PCW and older), then washed twice again in PBS, all at room temperature. Following dehydration in 70% and 100% ethanol for 2 minutes each, a 9 mm diameter (50 μl volume) SecureSeal hybridisation chamber (Grace Bio-Labs GBL621505-20EA) was adhered to each slide and used to hold subsequent reaction mixtures. Following rehydration in buffer WB3, probe hybridisation in buffer RM1 was conducted for 16 hours at 37°C. The 158-plex probe panel included 5 padlock probes per gene, the sequences of which are proprietary (10X Genomics CARTANA). The section was washed with PBS-T (PBS with 0.05% Tween-20) twice, then with buffer WB4 for 30 minutes at 37°C, and thrice again with PBS-T. Probe ligation in RM2 was conducted for 2 hours at 37°C and the section washed thrice with PBS-T, then rolling circle amplification in RM3 was conducted for 18 hours at 30°C. Following PBS-T washes, all rolling circle products (RCPs) were hybridised with LM (Cy5 labelling mix with DAPI) for 30 minutes at room temperature, the section was washed with PBS-T and dehydrated with 70% and 100% ethanol. The hybridisation chamber was removed and the slide mounted with SlowFade Gold Antifade Mountant (Thermo S36937). Imaging of Cy5-labelled RCPs at this stage acted as a QC step to confirm RCP (‘anchor’) generation and served to identify spots during decoding. Imaging was conducted using a Perkin Elmer Opera Phenix Plus High-Content Screening System in confocal mode with 1 μm z-step size, using a 63X (NA 1.15, 0.097 μm/pixel) water-immersion objective. Channels: DAPI (excitation 375 nm, emission 435-480 nm), Atto 425 (ex. 425 nm, em. 463-501 nm), Alexa Fluor 488 (ex. 488 nm, em. 500-550 nm), Cy3 (ex. 561 nm, em. 570-630 nm), Cy5 (ex. 640 nm, em. 650-760 nm).

Following imaging, each slide was de-coverslipped vertically in PBS (gently, with minimal agitation such that the coverslip ‘fell’ off to prevent damage to the tissue). The section was dehydrated with 70% and 100% ethanol, and a new hybridisation chamber secured to the slide. The previous cycle was stripped using 100% formamide (Thermo AM9342), which was applied fresh each minute for 5 minutes, then washed with PBS-T. Barcode labelling was conducted using two rounds of hybridisation, first an adapter probe pool (AP mixes AP1-AP6, in subsequent cycles), then a sequencing pool (SP mix, customised with Atto 425), each for 1 hour at 37°C with PBS-T washes in between and after. The section was dehydrated, the chamber removed, and the slide mounted and imaged as previously. This was repeated another five times to generate the full dataset of 7 cycles (anchor and 6 barcode bits).

### Multiplexed smFISH

Cryosections were processed using a Leica BOND RX to automate staining with the RNAscope Multiplex Fluorescent Reagent Kit v2 Assay (Advanced Cell Diagnostics, Bio-Techne), according to the manufacturers’ instructions. Probes may be found in Supplementary Table 7. Prior to staining, fresh frozen sections were post-fixed in 4% paraformaldehyde in PBS for 45 minutes at 4°C, then dehydrated through a series of 50%, 70%, 100%, and 100% ethanol, for 5 minutes each. Following manual pre-treatment, automated processing included digestion with Protease III for 15 minutes prior to probe hybridisation. Tyramide signal amplification with Opal 520, Opal 570, and Opal 650 (Akoya Biosciences) and TSA-biotin (TSA Plus Biotin Kit, Perkin Elmer) and streptavidin-conjugated Atto 425 (Sigma Aldrich) was used to develop RNAscope probe channels. Stained sections were imaged as for ISS above.

### Image-based in situ sequencing (ISS) decoding

We employed the ISS decoding pipeline outlined in Li et al ^114^. This pipeline consists of five distinct steps. Firstly, we performed image stitching using Acapella scripts provided by Perkin Elmer, which generated two-dimensional maximum intensity projections of all channels for each cycle. Next, we employed Microaligner^115^ to register all cycles based on DAPI signals using the default parameters. For cell segmentation, we utilised a scalable algorithm that leverages CellPose^116^ as the segmentation method. The expected cell size is set to 70 pixels in diameter and further expanded 10 pixels to mimic the cytoplasm. To decode the RNA molecules, we employed the PoSTcode algorithm^117^ with the following parameters: rna_spot_size=5, prob_threshold=0.6, trackpy_percentile=90, trackpy_separation=2. Furthermore, we assigned the decoded RNA molecules to segmented cells using STRtree and subsequently generated AnnData objects following the approach described by Virshup et al. ^118^. Finally, only cells with more than 4 RNA molecules were kept for downstream analysis.

### Visium processing and library preparation

Visium CytAssist Spatial Gene Expression for Fresh Frozen (10x Genomics) was performed following the manufacturer’s protocol. Regions of interest (ROIs) were selected based on the presence of microenvironments of bone-formation relevant to the droplet data (e.g. coronal suture) and aligned to the CytAssist machine gasket accordingly. Images were captured using a Hamamatsu S60 slide scanner at 40x magnification prior to conducting the Visium CytAssist protocol for subsequent alignment. Libraries were mapped with SpaceRanger (10x Genomics).

### Single-nuclei isolation and library preparation

Single-nuclei were isolated from fresh frozen samples through cryosectioning followed by mechanical dissociation as described in previous work ^119^. Briefly, 10 μm sections were homogenised in homogenization buffer (250 mM sucrose, 25 mM KCl, 5 mM MgCl_2_, 10 mM Tris-HCl, 1 mM dithiothreitol (DTT), 1× protease inhibitor, 0.4 U μl^−1^ RNaseIn, 0.2 U μl^−1^ SUPERaseIn and 0.1% Triton X-100 in nuclease-free water) using a glass Dounce tissue grinder set (Merck). Samples were dissociated with 10-20 strokes of a loose pestle ‘A’ followed by 10 strokes of a tight pestle ‘B’ when tissue fragments remained . The resulting mixture was passed through a 50 μm cell strainer, followed by centrifugation (500 g, 5 mins) the pellet was then resuspended in 300 μl of storage buffer (1× PBS, 4% BSA and 0.2 U μl^−1^ Protector RNaseIn) and passed through a 20 μm cell strainer. Nuclei were visualised and assessed for viability under microscopy following staining with trypan blue solution and were further processed for 10x Genomics Chromium Single Cell Multiome ATAC + Gene Expression according to the manufacturer’s protocol. Nuclei suspensions were loaded with a targeted nuclei recovery of 10,000 droplets per reaction. For some of the nuclei samples, mixtures of different donors were pooled within one reaction and later demultiplexed by genotype. Quality control of cDNA and final libraries was done using Bioanalyzer High Sensitivity DNA Analysis (Agilent). Libraries were sequenced using a NovaSeq 6000 (Illumina) with a minimum sequencing depth of 20,000 read pairs per droplet.

### Data preprocessing

Sequencing data were aligned to the human reference genome (GRCh38-2020-A-2.0.0) using CellRanger-ARC software (v.2.0.0). The called barcodes from 10x Multiome lanes with pooled genotypes from multiple donors were demultiplexed per Genotype using BAM outputs through Souporcell (v2.0) ^120^. Subsequently, the Souporcell outputs were clustered by genotype for metadata assignment to each barcode. Visium data were mapped to SpaceRanger (v. 1.1.0) using default input settings, and low-res Cytassist images were aligned to hi-res microscopy images of the processed slides using 10x Genomics LoupeBrowser (v 7.0) according to capture frame marker regions. For gene expression data, SoupX ^121^ was applied to remove background ambient RNA. For cellranger-arc called matrices that contained >16,000 droplets (exceeding the number expected from targeted droplet recovery) we increased the estimated global rho value by 0.1 to account for the potential of additional ambient RNA. Scrublet ^122^ was applied to estimate doublet probability and a score of >0.3 was used as a cutoff value. Droplets were filtered for >200 genes, and <5% mitochondrial and ribosomal reads.

For scATAC-seq, we applied ArchR^123^ (v1.0.2) to process the outputs from Cellranger-atac. Initial per-cell quality control was performed considering the number of unique nuclear fragments, signal-to-background ratio and the fragment size distribution. Moreover, cells with TSSenrichment score<7 and nFrags< 1000 were removed. Doublets were discarded using the default settings. Initial clustering was performed at resolution = 0.2 with top 40 dimensions from Iterative Latent Semantic Indexing (LSI). Then, pseudo-bulk replicates were made for each broad cell type per region from the initial clustering results. Peak calling (501 bp fixed-width peaks) was performed based on pseudo-bulk coverages by macs2. Then, a cell-by-peak count matrix was obtained and exported. We applied muon^124^ (0.1.2) for normalization, LSI dimension reduction, and clustering analysis with BBKNN^125^ to correct the batch effects from anatomical regions and donors for obtaining ATAC embedding. Gene scores based on chromatin accessibility around gene bodies were calculated. Moreover, we applied MultiVI^126^ to get a joint embedding for snRNA-seq and scATAC-seq.

### Doublet detection

To interrogate doublet capture, we conducted an adapted scrublet workflow that was previously described^127^. Briefly, per-droplet scrublet scores were first determined for cellranger-arc count-matrices from each 10x multiome (gene-expression) lane independently. The droplets were then overclustered through the standard scanpy workflow using default parameters up to leiden clustering. Each individual cluster was further clustered. A per-cluster median of scrublet scores were computed. A normal distribution of doublet score, centred at the score median with a standard deviation estimated from the median absolute deviation (MAD) was used to compute p-values for each of the clusters. After false-discovery rate adjustment using benjamini-hochberg correction, a p-value of >0.6 was deemed as a cutoff value of good quality cells as doublets are significant outliers.

### Cell cluster annotation

We adopted a hierarchical clustering approach by first conducting Leiden clustering on the global integrated scVI (hidden layers=256, latent variables=52, dispersion=‘gene-batch’) RNA embeddings to obtain broad clusters. To validate these we utilised Celltypist to train a model on cell states in the embryonic limb bud ^11,128,129^, and transferred labels onto our embedding for inspection. We utilise this information in addition to canonical marker genes, to annotate broad clusters and subset sub-lineages. For sub-lineages e.g. (Chondrocytes, Fibroblasts, Bone-related populations, Schwann Cells, Immune cells, Endothelial cells), we further embedded these using scVI (hidden layers=256, latent variables=52, dispersion=‘gene-batch’) and conducted Leiden clustering (resolution = 0.6), followed by DEG analyses (method=’wilcoxon’) to obtain cluster markers. We additionally utilised the inferred spatial location of cell states (described below) to inform annotations.

### Spatial mapping using Cell2location

We performed Cell2location for deconvolution of visium cytassist using annotated snRNA-seq data from skull base and calvaira cell states as input. The donor was used as batch variables, and the libraries were considered as covariates in the regression model. For spatial mapping, we estimated 30 cells per voxel based on histological data, and hyperparameter detection_alpha of 20 were applied for per-voxel normalisation.

### ISS Patcher

ISS_patcher is a simple package for approximating features not experimentally captured in low-dimensional data based on related, high-dimensional data. It was developed as an approach to approximate expression signatures for genes missing in ISS data using matched snRNA-Seq data as a reference in this study. First, a shared feature space between both datasets was identified by subsetting the 155-158 genes present in the ISS pool, followed by separate normalisation to median total cell counts, log-transformation and z-scoring for both modalities. Then, the 15 nearest neighbours in scRNA-Seq space were identified for each ISS cell with the Annoy python package, and the genes absent from ISS were imputed as the average raw counts of the scRNA-Seq neighbours.

### Visium axis annotation using OrganAxis

Our Visium cranium sample was annotated with TissueTag by semi-automatic mode to generate a one-dimensional maturation axis. Regions of the developing bone were first manually annotated based on H&E features. Tissue regions that did not include bone-forming niches were excluded from annotation. The annotation categories that were stored included multiple regions of the coronal suture (level_0 to level_2 annotation), stemming from the central-most portion, an osteogenic front (level_3 annotation) harbouring histological features of osteoprogenitors and osteogenic zones (level_4 to level_7 annotation) from the emergence of histological osteoblasts. All annotations were saved as TissueTag output format, which logs the annotation resolution, the pixels per μm (ppm) as well as pixel value interpretation of annotation names (e.g. 0 = “Suture”) and colours (e.g. “Osteogenic Front”: “red”). To robustly and efficiently migrate TissueTag annotations to the Visium objects, we first transferred TissueTag annotations from pixel space to a high-resolution hexagonal grid space (15 µm spot diameter and 15 µm point-to-point centre distance with no gap between spots) based on the median pixel value of the centre of the spot (radius/4) in the annotated image. Next, to generate continuous annotations for Visium data we measured for each spot in the hexagonal high-resolution grid the mean Euclidean distance to the 10 nearest points from each annotated structure in the level_0 annotation as well as the distance from the closest point for structures in annotation_level_1. All annotations were mapped to the Visium spots by proximity of the spot annotation grid to the nearest corresponding spot in the Visium array.

### Gene regulatory network analysis

The SCENIC+ ^130^ pipeline was used to predict transcription factors and putative target genes as well as regulatory genomic regions harbouring binding sites. The fragment matrix of peaks called with macs2 and processed within ArchR ^123^ together with the corresponding RNA count matrix were used as inputs. Meta-cells were created by clustering cells into groups of around 10-15 cells based on their RNA profiles and subsequent aggregation of counts and fragments. The pipeline was applied to subsets of the dataset corresponding to individual lineages: First, CisTopic was applied to identify region topics and differentially accessible regions from the fragment counts as candidate regions for TF binding. CisTarget was then run to scan the regions for transcription factor binding sites and GRNBoost2 ^131^ was used to link TFs and regions to target genes based on co-expression/accessibility. Enriched TF motifs in the regions linked to target genes were used to construct TF-region and TF-gene regulons. Finally, regulon activity scores (AUC) were computed with AUCell based on target-gene expression and target region accessibility, and regulon specificity scores (RSS) derived from them. Networks of TFs, regions and target genes (eGRNs) were constructed by linking individual regulons. TF-enhancer-gene links for all subsets (osteogenesis, chondrogenesis, fibrogenesis, early joint progenitors, immune, schwann) can be found in Supplementary Table 9.

### Trajectory analysis

For trajectory construction in the osteogenic subcompartment, non-cycling droplets were subsetted and X_scVI were used as projections for palantir in order to obtain multiuscale diffusion space. A neighbourhood graph was generated on the diffusion space using sc.pp.neighbors, and the first two PCs were used as initial positions to create ForceAtlas2 embeddings using sc.tl.draw_graph. The FA embeddings were exported into R, and monocle3 ^132,133^ was used to find a principal graph and define pseudotime. Differentially expressed genes were then computed along pseudotime using a graph-based test (morans’ I) ^134,135^ and the principal graph in monocle3, which allows identification of genes upregulated at any point in pseudotime. The results were visualised with heatmaps using the complexHeatmap ^136^ and seriation ^137^ packages, after smoothing gene expression with smoothing splines in R (smooth.spline, df=12). Velocity analysis^138^ was performed using scvelo^139^ version 0.2.3. Spliced and unspliced read counts were computed with velocyto from the unprocessed data, before using scvelo.pp.moments, scvelo.tl.velocity and scvelo.tl.velocity_graph to compute velocities for the preprocessed droplets.

### *In silico* TF perturbations

CellOracle ^140^ was used with the osteogenesis trajectory created with scFates ^141^ and the regulons predicted with SCENIC+ ^130^ for the same cells were imported into CellOracle as a base GRN. Cells were aggregated into meta-cells of 10-15 cells and linear models explaining TF- from target gene expression were fitted with CellOracle per cell cluster. Regulon-based TF perturbation vectors were inferred using the cell cluster-specific models to predict effects of TF overexpression and knockout. Diffusion pseudotime ^142^ was then computed for IM and EC lineages separately by selecting corresponding starting points. The pseudotime gradients were used to derive pseudotime-based differentiation vectors, and the pseudotime-perturbation vector cross-product was computed to obtain perturbation scores (PS). These PS indicate, whether the *in silico* perturbation of a TF is consistent with or opposes differentiation along a lineage (osteogenesis). The simulations were carried out systematically, overexpressing and knocking out all TFs in the GRN. For each TF and condition the PS were then averaged per cell cluster and summarised in a table to screen for TFs promoting or inhibiting osteogenesis.

### fGWAS

Functional GWAS analysis (fGWAS) ^143^ was applied to identify disease relevant cell clusters as described in detail in ^144^ (https://github.com/natsuhiko/PHM). The model makes use of full summary statistics from GWAS studies, linking SNPs to genes and captures a general trend between gene expression and disease association of linked loci for each cell cluster. At the same time, it also corrects for linkage disequilibrium (LD) and other relevant factors. We used full GWAS summary statistics obtained from the EBI GWAS catalog, open targets and knee and hip osteoarthritis (OA) as well as total knee (TKR) and hip (THR) replacement from ^145^ (Supplementary Table 8).

### SNP2Cell

We used a network propagation ^91^ approach to integrate GWAS summary statistics and cell cluster marker gene based scores for prioritising disease relevant and cell cluster specific subunits of our TF-network. First, scores per SNP were computed from downloaded summary statistics and weighted by linkage-disequilibrium. Then the scores were mapped to a gene regulatory network, here an eGRN computed with SCENIC+ for the corresponding lineage. Since the used networks contain TFs and target genes, and also regions with TF-binding sites as nodes, SNP scores were mapped both to genes and regions, representing distal regulatory elements. The scores were then propagated across the network using a random walk with restart (or personalised page-rank) process. This integrates the contribution of individual SNPs, with signals converging around relevant network nodes. The procedure was repeated with 1000 randomly permuted scores to compute permutation-test results and z-scores. Next, differential expression based marker gene scores for each cell cluster were propagated in the same way, resulting in cell-cluster specificity scores for each network node. The SNP and expression based scores were then combined per cell cluster as in ^146^ by using the minimum for each node. The final scores were thresholded and the resulting connected components obtained as enriched sub-networks. The method has been compiled into a tool we called SNP2CELL and is available at (https://github.com/Teichlab/snp2cell).

### Cell-cell interactions

Ligand-receptor interactions were inferred using ‘cpdb_analysis_method.call’ in CellPhoneDB (v4.0.0). We only considered genes with more than 10% of cells demonstrating expression within each cell cluster considered and interactions with P >0.001 were removed. We used NicheNet (v1.1.1) to identify different interactions between endochondral ossification and intramembranous niches. We first calculate differentially expressed genes (DEGs) of osteoblastic cell clusters and tip cells between two niches using standard Seurat Wilcoxon test and min LFC per cell cluster were used to summarise the differentially expressed ligands and receptors. Then, top 1000 DEGs were applied for calculating the ligand activities. We prioritised the ligand-receptor links with default settings. Top 10 ligands and their top scoring receptors were visualised in heatmaps.

## References

1. O’Rahilly, R. & Gardner, E. The initial appearance of ossification in staged human embryos. Am. J. Anat. 134, 291–307 (1972).

2. Faro, C., Benoit, B., Wegrzyn, P., Chaoui, R. & Nicolaides, K. H. Three-dimensional sonographic description of the fetal frontal bones and metopic suture. Ultrasound Obstet. Gynecol. 26, 618–621 (2005).

3. Edwards, J. C. et al. The formation of human synovial joint cavities: a possible role for hyaluronan and CD44 in altered interzone cohesion. J. Anat. 185, 355 (1994).

4. Pazzaglia, U. E. et al. Long bone human anlage longitudinal and circumferential growth in the fetal period and comparison with the growth plate cartilage of the postnatal age. Microsc. Res. Tech. 82, 190–198 (2019).

5. Pazzaglia, U. E. et al. Study of Endochondral Ossification in Human Fetalcartilage Anlagen of Metacarpals: Comparative Morphology of Mineral Deposition in Cartilage and in the Periosteal Bone Matrix. Anat. Rec. 301, 571–580 (2018).

6. Wilkie, A. O. M., Johnson, D. & Wall, S. A. Clinical genetics of craniosynostosis. Curr. Opin. Pediatr. 29, 622 (2017).

7. Wilkie, A. O. M. Craniosynostosis: Genes and Mechanisms. Hum. Mol. Genet. 6, 1647–1656 (1997).

8. Twigg, S. R. & Wilkie, A. O. A Genetic-Pathophysiological Framework for Craniosynostosis. Am. J. Hum. Genet. 97, (2015).

9. Rice, S. J. et al. Genetic risk of osteoarthritis operates during human skeletogenesis. Hum. Mol. Genet. 32, (2023).

10. Capellini, T. D. et al. Ancient selection for derived alleles at a GDF5 enhancer influencing human growth and osteoarthritis risk. Nat. Genet. 49, (2017).

11. Zhang, B., et al. A human embryonic limb cell atlas resolved in space and time. *bioRxiv* 2022.04.27.489800 (2023) doi:10.1101/2022.04.27.489800.

12. Jardine, L. et al. Blood and immune development in human fetal bone marrow and Down syndrome. Nature 598, 327–331 (2021).

13. He, P. et al. The changing mouse embryo transcriptome at whole tissue and single-cell resolution. Nature 583, 760–767 (2020).

14. Pdgfra regulates multipotent cell differentiation towards chondrocytes via inhibiting Wnt9a/beta-catenin pathway during chondrocranial cartilage development. Dev. Biol. 466, 36–46 (2020).

15. Xi, H. et al. A Human Skeletal Muscle Atlas Identifies the Trajectories of Stem and Progenitor Cells across Development and from Human Pluripotent Stem Cells. Cell Stem Cell 27, 181–185 (2020).

16. Hita-Contreras, F., et al. Development of the human shoulder joint during the embryonic and early fetal stages: anatomical considerations for clinical practice. J. Anat. 232, 422 (2018).

17. Feng, C. et al. Lgr5 and Col22a1 Mark Progenitor Cells in the Lineage toward Juvenile Articular Chondrocytes. Stem cell reports 13, (2019).

18. Shwartz, Y., Viukov, S., Krief, S. & Zelzer, E. Joint Development Involves a Continuous Influx of Gdf5-Positive Cells. Cell Rep. 15, (2016).

19. Bian, Q. et al. A single cell transcriptional atlas of early synovial joint development. Development 147, (2020).

20. Lee, W. et al. Synergy between Piezo1 and Piezo2 channels confers high-strain mechanosensitivity to articular cartilage. Proc. Natl. Acad. Sci. U. S. A. 111, (2014).

21. Beccari, L. et al. Dbx2 regulation in limbs suggests interTAD sharing of enhancers. Dev. Dyn. 250, 1280 (2021).

22. Chen, H. et al. Runx2 Regulates Endochondral Ossification through Control of Chondrocyte Proliferation and Differentiation. J. Bone Miner. Res. 29, 2653 (2014).

23. Runx2 is required for hypertrophic chondrocyte mediated degradation of cartilage matrix during endochondral ossification. Matrix Biology Plus 12, 100088 (2021).

24. Ono, K. et al. Dmrt2 promotes transition of endochondral bone formation by linking Sox9 and Runx2. Communications Biology 4, 1–13 (2021).

25. Snelling, S. J., Hulley, P. A. & Loughlin, J. BMP5 activates multiple signaling pathways and promotes chondrogenic differentiation in the ATDC5 growth plate model. Growth Factors 28, (2010).

26. Craft, A. M. et al. Specification of chondrocytes and cartilage tissues from embryonic stem cells. Development 140, 2597–2610 (2013).

27. Pothiawala, A. et al. GDF5+ chondroprogenitors derived from human pluripotent stem cells preferentially form permanent chondrocytes. Development 149, dev196220 (2022).

28. Sugimoto, Y. et al. Scx+/Sox9+ progenitors contribute to the establishment of the junction between cartilage and tendon/ligament. Development 140, 2280–2288 (2013).

29. Arostegui, M., Scott, R. W., Böse, K. & Underhill, T. M. Cellular taxonomy of Hic1+ mesenchymal progenitor derivatives in the limb: from embryo to adult. Nat. Commun. 13, 1–20 (2022).

30. Buechler, M. B. et al. Cross-tissue organization of the fibroblast lineage. Nature 593, 575–579 (2021).

31. Palumbo-Zerr, K. et al. Composition of TWIST1 dimers regulates fibroblast activation and tissue fibrosis. Ann. Rheum. Dis. 76, 244–251 (2017).

32. Twist2-driven chromatin remodeling governs the postnatal maturation of dermal fibroblasts. Cell Rep. 39, 110821 (2022).

33. Mascharak, S. et al. Preventing Engrailed-1 activation in fibroblasts yields wound regeneration without scarring. Science 372, (2021).

34. Sontake, V., et al. Wilms’ tumor 1 drives fibroproliferation and myofibroblast transformation in severe fibrotic lung disease. JCI Insight 3, (2018).

35. Shi, Y. et al. Transcription Factor SOX5 Promotes the Migration and Invasion of Fibroblast-Like Synoviocytes in Part by Regulating MMP-9 Expression in Collagen-Induced Arthritis. Front. Immunol. 9, (2018).

36. Foxc1 promotes the proliferation of fibroblast-like synoviocytes in rheumatoid arthritis via PI3K/AKT signalling pathway. Tissue and Cell 53, 15–22 (2018).

37. Wang, J. et al. Forkhead box C1 promotes the pathology of osteoarthritis by upregulating β-catenin in synovial fibroblasts. FEBS J. 287, (2020).

38. Bobowski-Gerard, M. et al. Functional genomics uncovers the transcription factor BNC2 as required for myofibroblastic activation in fibrosis. Nat. Commun. 13, 1–20 (2022).

39. Morriss-Kay, G. M. & Wilkie, A. O. M. Growth of the normal skull vault and its alteration in craniosynostosis: insights from human genetics and experimental studies. J. Anat. 207, 637–653 (2005).

40. Zhao, H. et al. The suture provides a niche for mesenchymal stem cells of craniofacial bones. Nat. Cell Biol. 17, 386–396 (2015).

41. Zhao, Q., Behringer, R. R. & de Crombrugghe, B. Prenatal folic acid treatment suppresses acrania and meroanencephaly in mice mutant for the Cart1 homeobox gene. Nat. Genet. 13, 275–283 (1996).

42. Pini, J. et al. ALX1-related frontonasal dysplasia results from defective neural crest cell development and migration. EMBO Mol. Med. 12, (2020).

43. Eze, U. C., Bhaduri, A., Haeussler, M., Nowakowski, T. J. & Kriegstein, A. R. Single-cell atlas of early human brain development highlights heterogeneity of human neuroepithelial cells and early radial glia. Nat. Neurosci. 24, 584–594 (2021).

44. Rocha, M. et al. From head to tail: regionalization of the neural crest. Development 147, dev193888 (2020).

45. Farmer, D. T. et al. The developing mouse coronal suture at single-cell resolution. Nat. Commun. 12, (2021).

46. Debnath, S. et al. Discovery of a periosteal stem cell mediating intramembranous bone formation. Nature 562, 133–139 (2018).

47. Cesario, J. M. et al. Anti-osteogenic function of a LIM-homeodomain transcription factor LMX1B is essential to early patterning of the calvaria. Dev. Biol. 443, 103 (2018).

48. Zhang, Q. et al. RUNX2 co-operates with EGR1 to regulate osteogenic differentiation through Htra1 enhancers. J. Cell. Physiol. 235, (2020).

49. Zaman, G. et al. Loading-related Regulation of Transcription Factor EGR2/Krox-20 in Bone Cells Is ERK1/2 Protein-mediated and Prostaglandin, Wnt Signaling Pathway-, and Insulin-like Growth Factor-I Axis-dependent. J. Biol. Chem. 287, 3946 (2012).

50. Rached, M. T. et al. FoxO1 is a positive regulator of bone formation by favoring protein synthesis and resistance to oxidative stress in osteoblasts. Cell Metab. 11, (2010).

51. Zalc, A., Rattenbach, R., Auradé, F., Cadot, B. & Relaix, F. Pax3 and Pax7 play essential safeguard functions against environmental stress-induced birth defects. Dev. Cell 33, (2015).

52. Ono, N., Ono, W., Nagasawa, T. & Kronenberg, H. M. A subset of chondrogenic cells provides early mesenchymal progenitors in growing bones. Nat. Cell Biol. 16, 1157–1167 (2014).

53. Peters, H., Neubüser, A., Kratochwil, K. & Balling, R. Pax9-deficient mice lack pharyngeal pouch derivatives and teeth and exhibit craniofacial and limb abnormalities. Genes Dev. 12, (1998).

54. Holmes, G. et al. Single-cell analysis identifies a key role for Hhip in murine coronal suture development. Nat. Commun. 12, 1–16 (2021).

55. Orestes-Cardoso, S. M. et al. Postnatal Msx1 expression pattern in craniofacial, axial, and appendicular skeleton of transgenic mice from the first week until the second year. Dev. Dyn. 221, 1–13 (2001).

56. Zanotti, S. & Canalis, E. Activation of Nfatc2 in osteoblasts causes osteopenia. J. Cell. Physiol. 230, (2015).

57. Ranger, A. M. et al. The Nuclear Factor of Activated T Cells (Nfat) Transcription Factor Nfatp (Nfatc2) Is a Repressor of Chondrogenesis. J. Exp. Med. 191, 9–22 (2000).

58. Loss of Iroquois homeobox transcription factors 3 and 5 in osteoblasts disrupts cranial mineralization. Bone Reports 5, 86–95 (2016).

59. Maes, C. et al. Osteoblast Precursors, but Not Mature Osteoblasts, Move into Developing and Fractured Bones along with Invading Blood Vessels. Dev. Cell 19, 329–344 (2010).

60. Percival, C. J. & Richtsmeier, J. T. Angiogenesis and intramembranous osteogenesis. Dev. Dyn. 242, 909–922 (2013).

61. Zhu, K. et al. ATF4 promotes bone angiogenesis by increasing VEGF expression and release in the bone environment. J. Bone Miner. Res. 28, 1870 (2013).

62. Kronenberg, H. M. Developmental regulation of the growth plate. Nature 423, 332–336 (2003).

63. Kusumbe, A. P., Ramasamy, S. K. & Adams, R. H. Coupling of angiogenesis and osteogenesis by a specific vessel subtype in bone. Nature 507, 323–328 (2014).

64. Wang, Y. et al. Ephrin-B2 controls VEGF-induced angiogenesis and lymphangiogenesis. Nature 465, (2010).

65. Wang, F. et al. RACK1 regulates VEGF/Flt1-mediated cell migration via activation of a PI3K/Akt pathway. J. Biol. Chem. 286, (2011).

66. Ruehle, M. A. et al. Mechanical Regulation of Microvascular Angiogenesis. *bioRxiv* 2020.01.14.906354 (2020) doi:10.1101/2020.01.14.906354.

67. Efremova, M., Vento-Tormo, M., Teichmann, S. A. & Vento-Tormo, R. CellPhoneDB: inferring cell-cell communication from combined expression of multi-subunit ligand-receptor complexes. Nat. Protoc. 15, 1484–1506 (2020).

68. Ramasamy, S. K., Kusumbe, A. P., Wang, L. & Adams, R. H. Endothelial Notch activity promotes angiogenesis and osteogenesis in bone. Nature 507, 376–380 (2014).

69. Miraoui, H. et al. Fibroblast growth factor receptor 2 promotes osteogenic differentiation in mesenchymal cells via ERK1/2 and protein kinase C signaling. J. Biol. Chem. 284, (2009).

70. Kevorkova, O. et al. Low-Bone-Mass Phenotype of Deficient Mice for the Cluster of Differentiation 36 (CD36). PLoS One 8, e77701 (2013).

71. Baron, R. & Kneissel, M. WNT signaling in bone homeostasis and disease: from human mutations to treatments. Nat. Med. 19, 179–192 (2013).

72. Yu, X., Huang, Y., Collin-Osdoby, P. & Osdoby, P. CCR1 Chemokines Promote the Chemotactic Recruitment, RANKL Development, and Motility of Osteoclasts and Are Induced by Inflammatory Cytokines in Osteoblasts. J. Bone Miner. Res. 19, 2065–2077 (2004).

73. DPP-4 inhibitor impedes lipopolysaccharide-induced osteoclast formation and bone resorption in vivo. Biomed. Pharmacother. 109, 242–253 (2019).

74. Tomlinson, R. E. et al. NGF-TrkA Signaling by Sensory Nerves Coordinates the Vascularization and Ossification of Developing Endochondral Bone. Cell Rep. 16, (2016).

75. Hu, B. et al. Sensory nerves regulate mesenchymal stromal cell lineage commitment by tuning sympathetic tones. J. Clin. Invest. 130, (2020).

76. Tower, R. J. et al. Spatial transcriptomics reveals a role for sensory nerves in preserving cranial suture patency through modulation of BMP/TGF-β signaling. Proc. Natl. Acad. Sci. U. S. A. 118, (2021).

77. Shum, L. & Nuckolls, G. The life cycle of chondrocytes in the developing skeleton. Arthritis Res. 4, 94–106 (2002).

78. Taïhi, I., Nassif, A., Isaac, J., Fournier, B. P. & Ferré, F. Head to Knee: Cranial Neural Crest-Derived Cells as Promising Candidates for Human Cartilage Repair. Stem Cells Int. 2019, 9310318 (2019).

79. Chilton, J. K. & Guthrie, S. Cranial expression of class 3 secreted semaphorins and their neuropilin receptors. Dev. Dyn. 228, 726–733 (2003).

80. Berndt, J. D. & Halloran, M. C. Semaphorin 3d promotes cell proliferation and neural crest cell development downstream of TCF in the zebrafish hindbrain. Development 133, 3983–3992 (2006).

81. DeTomaso, D. & Yosef, N. Hotspot identifies informative gene modules across modalities of single-cell genomics. Cell Syst. 12, 446–456.e9 (2021).

82. Wang, Z., Liu, B., Lin, K., Duan, C. & Wang, C. The presence and degradation of nerve fibers in articular cartilage of neonatal rats. J. Orthop. Surg. Res. 17, 331 (2022).

83. Yueh, Y. G., Gardner, D. P. & Kappen, C. Evidence for regulation of cartilage differentiation by the homeobox gene Hoxc-8. Proc. Natl. Acad. Sci. U. S. A. 95, (1998).

84. Cartilage repair using bone morphogenetic protein 4 and muscle-derived stem cells. doi:10.1002/art.21632.

85. Yin, Z. et al. Atlas of Musculoskeletal Stem Cells with the Soft and Hard Tissue Differentiation Architecture. Adv. Sci. Lett. 7, (2020).

86. Grimaldi, A., Comai, G., Mella, S. & Tajbakhsh, S. Identification of bipotent progenitors that give rise to myogenic and connective tissues in mouse. Elife 11, (2022).

87. He, J. et al. Dissecting human embryonic skeletal stem cell ontogeny by single-cell transcriptomic and functional analyses. Cell Res. 31, 742–757 (2021).

88. Xie, M. et al. Schwann cell precursors contribute to skeletal formation during embryonic development in mice and zebrafish. Proc. Natl. Acad. Sci. U. S. A. 116, 15068–15073 (2019).

89. Richard, D. et al. Evolutionary Selection and Constraint on Human Knee Chondrocyte Regulation Impacts Osteoarthritis Risk. Cell 181, 362–381.e28 (2020).

90. Wilkinson, J. M. & Zeggini, E. The Genetic Epidemiology of Joint Shape and the Development of Osteoarthritis. Calcif. Tissue Int. 109, 257–276 (2021).

91. Cowen, L., Ideker, T., Raphael, B. J. & Sharan, R. Network propagation: a universal amplifier of genetic associations. Nat. Rev. Genet. 18, 551–562 (2017).

92. Heinonen, J. et al. Defects in chondrocyte maturation and secondary ossification in mouse knee joint epiphyses due to Snorc deficiency. Osteoarthritis Cartilage 25, 1132–1142 (2017).

93. Lee, H.-S. et al. Activation of Integrin-RACK1/PKCalpha signalling in human articular chondrocyte mechanotransduction. Osteoarthritis Cartilage 10, 890–897 (2002).

94. Zhang, F. et al. NFATc1 marks articular cartilage progenitors and negatively determines articular chondrocyte differentiation. Elife 12, (2023).

95. Greenblatt, M. B. et al. NFATc1 and NFATc2 repress spontaneous osteoarthritis. Proc. Natl. Acad. Sci. U. S. A. 110, 19914–19919 (2013).

96. Ren, R. et al. The role of Ca2+/Calcineurin/NFAT signalling pathway in osteoblastogenesis. Cell Prolif. 54, e13122 (2021).

97. Genetics of developmental dysplasia of the hip. Eur. J. Med. Genet. 63, 103990 (2020).

98. Zhang, X. et al. Lipid peroxidation in osteoarthritis: focusing on 4-hydroxynonenal, malondialdehyde, and ferroptosis. Cell Death Discovery 9, 1–13 (2023).

99. Mathijssen, I. M. et al. Tracing craniosynostosis to its developmental stage through bone center displacement. J. Craniofac. Genet. Dev. Biol. 19, (1999).

100. Lajeunie, E., Le Merrer, M., Bonaïti-Pellie, C., Marchac, D. & Renier, D. Genetic study of nonsyndromic coronal craniosynostosis. Am. J. Med. Genet. 55, (1995).

101. Wilkie, A. O. et al. Prevalence and complications of single-gene and chromosomal disorders in craniosynostosis. Pediatrics 126, (2010).

102. Sharma, V. P. et al. Mutations in TCF12, encoding a basic helix-loop-helix partner of TWIST1, are a frequent cause of coronal craniosynostosis. Nat. Genet. 45, 304–307 (2013).

103. Yoon, W. J. et al. The Boston-type craniosynostosis mutation MSX2 (P148H) results in enhanced susceptibility of MSX2 to ubiquitin-dependent degradation. J. Biol. Chem. 283, (2008).

104. James, A. W., Levi, B., Xu, Y., Carre, A. L. & Longaker, M. T. Retinoic acid enhances osteogenesis in cranial suture-derived mesenchymal cells: potential mechanisms of retinoid-induced craniosynostosis. Plast. Reconstr. Surg. 125, 1352–1361 (2010).

105. Clouthier, D. E. et al. Cranial and cardiac neural crest defects in endothelin-A receptor-deficient mice. Development 125, 813–824 (1998).

106. Spence, S., Anderson, C., Cukierski, M. & Patrick, D. Teratogenic effects of the endothelin receptor antagonist L-753,037 in the rat. Reprod. Toxicol. 13, 15–29 (1999).

107. Bruderer, M., Richards, R. G., Alini, M. & Stoddart, M. J. Role and regulation of RUNX2 in osteogenesis. Eur. Cell. Mater. 28, 269–286 (2014).

108. van Lierop, A. H. et al. Van Buchem disease: clinical, biochemical, and densitometric features of patients and disease carriers. J. Bone Miner. Res. 28, 848–854 (2013).

109. Collette, N. M. et al. Targeted deletion of Sost distal enhancer increases bone formation and bone mass. Proc. Natl. Acad. Sci. U. S. A. 109, 14092–14097 (2012).

110. Liu, N. et al. DNA binding-dependent and -independent functions of the Hand2 transcription factor during mouse embryogenesis. Development 136, 933 (2009).

111. Xiong, W. et al. Hand2 is required in the epithelium for palatogenesis in mice. Dev. Biol. 330, 131 (2009).

112. Zhou, C. et al. Lhx8 Mediated Wnt and TGFβ Pathways in Tooth Development and Regeneration. Biomaterials 63, 35 (2015).

113. Lee, H., Marco Salas, S., Gyllborg, D. & Nilsson, M. Direct RNA targeted in situ sequencing for transcriptomic profiling in tissue. Sci. Rep. 12, 1–9 (2022).

114. Li, T. et al. WebAtlas pipeline for integrated single cell and spatial transcriptomic data. *bioRxiv* 2023.05.19.541329 (2023) doi:10.1101/2023.05.19.541329.

115. GitHub - VasylVaskivskyi/microaligner: Image registration (alignment) software for large microscopy images. GitHub https://github.com/VasylVaskivskyi/microaligner.

116. Pachitariu, M. & Stringer, C. Cellpose 2.0: how to train your own model. Nat. Methods 19, 1634–1641 (2022).

117. Gataric, M., et al. PoSTcode: Probabilistic image-based spatial transcriptomics decoder. *bioRxiv* (2021) doi:10.1101/2021.10.12.464086.

118. Virshup, I., Rybakov, S., Theis, F. J., Angerer, P. & Wolf, F. A. anndata: Annotated data. *bioRxiv* (2021) doi:10.1101/2021.12.16.473007.

119. Litviňuková, M. et al. Cells of the adult human heart. Nature 588, (2020).

120. Heaton, H. et al. Souporcell: robust clustering of single-cell RNA-seq data by genotype without reference genotypes. Nat. Methods 17, 615–620 (2020).

121. Young, M. D. & Behjati, S. SoupX removes ambient RNA contamination from droplet-based single-cell RNA sequencing data. Gigascience 9, giaa151 (2020).

122. Wolock, S. L., Lopez, R. & Klein, A. M. Scrublet: Computational Identification of Cell Doublets in Single-Cell Transcriptomic Data. Cell systems 8, (2019).

123. Granja, J. M. et al. ArchR is a scalable software package for integrative single-cell chromatin accessibility analysis. Nat. Genet. 53, 403–411 (2021).

124. Bredikhin, D., Kats, I. & Stegle, O. MUON: multimodal omics analysis framework. Genome Biol. 23, 1–12 (2022).

125. Polański, K. et al. BBKNN: fast batch alignment of single cell transcriptomes. Bioinformatics 36, (2020).

126. Ashuach, T. et al. MultiVI: deep generative model for the integration of multimodal data. Nat. Methods 20, 1222–1231 (2023).

127. Popescu, D.-M. et al. Decoding human fetal liver haematopoiesis. Nature 574, 365–371 (2019).

128. Xu, C. et al. Automatic cell type harmonization and integration across Human Cell Atlas datasets. *bioRxiv* 2023.05.01.538994 (2023) doi:10.1101/2023.05.01.538994.

129. Domínguez Conde, C., et al. Cross-tissue immune cell analysis reveals tissue-specific features in humans. Science 376, eabl5197 (2022).

130. Bravo González-Blas, C., et al. SCENIC+: single-cell multiomic inference of enhancers and gene regulatory networks. Nat. Methods 1–13 (2023).

131. Moerman, T. et al. GRNBoost2 and Arboreto: efficient and scalable inference of gene regulatory networks. Bioinformatics 35, 2159–2161 (2019).

132. Trapnell, C. et al. The dynamics and regulators of cell fate decisions are revealed by pseudotemporal ordering of single cells. Nat. Biotechnol. 32, 381–386 (2014).

133. Cao, J. et al. The single-cell transcriptional landscape of mammalian organogenesis. Nature vol. 566 496–502 Preprint at 10.1038/s41586-019-0969-x (2019).

134. Moran, P. A. P. Notes on continuous stochastic phenomena. Biometrika 37, 17–23 (1950).

135. Cao, J. et al. The single-cell transcriptional landscape of mammalian organogenesis. Nature vol. 566 496–502 Preprint at 10.1038/s41586-019-0969-x (2019).

136. Gu, Z., Eils, R. & Schlesner, M. Complex heatmaps reveal patterns and correlations in multidimensional genomic data. Bioinformatics vol. 32 2847–2849 Preprint at 10.1093/bioinformatics/btw313 (2016).

137. Hahsler, M., Hornik, K. & Buchta, C. Getting Things in Order: An Introduction to theRPackageseriation. Journal of Statistical Software vol. 25 Preprint at 10.18637/jss.v025.i03 (2008).

138. La Manno, G. et al. RNA velocity of single cells. Nature 560, 494–498 (2018).

139. Bergen, V., Lange, M., Peidli, S., Wolf, F. A. & Theis, F. J. Generalizing RNA velocity to transient cell states through dynamical modeling. Nat. Biotechnol. 38, 1408–1414 (2020).

140. Kamimoto, K. et al. Dissecting cell identity via network inference and in silico gene perturbation. Nature 614, 742–751 (2023).

141. Faure, L., Soldatov, R., Kharchenko, P. V. & Adameyko, I. scFates: a scalable python package for advanced pseudotime and bifurcation analysis from single-cell data. Bioinformatics 39, (2023).

142. Haghverdi, L., Büttner, M., Wolf, F. A., Buettner, F. & Theis, F. J. Diffusion pseudotime robustly reconstructs lineage branching. Nat. Methods 13, 845–848 (2016).

143. Pickrell, J. K. Joint analysis of functional genomic data and genome-wide association studies of 18 human traits. Am. J. Hum. Genet. 94, 559–573 (2014).

144. Elmentaite, R. et al. Cells of the human intestinal tract mapped across space and time. Nature 597, 250–255 (2021).

145. Boer, C. G. et al. Deciphering osteoarthritis genetics across 826,690 individuals from 9 populations. Cell 184, 6003–6005 (2021).

146. Paull, E. O. et al. Discovering causal pathways linking genomic events to transcriptional states using Tied Diffusion Through Interacting Events (TieDIE). Bioinformatics 29, 2757–2764 (2013).

