## Supplementary Information for "A multiomic atlas of human early skeletal development"

**Supplementary Table 1. Samples Overview**

List of donor ID obtained under approvals of REC 96/085 with corresponding anatomical regions, stage (PCW) and number of samples within each region processed across the skeleton. Stage was determined using crown-rump length (CRL) calculations: PCW (days) = 0.9022 × CRL (mm) + 27.372. (See SuppTable_1.xlsx)

**Supplementary Table 2. Cell state marker genes**

Differentially expressed genes (DEGs) for major transcriptomic clusters defined in this study across compartments. (See SuppTable_2.xlsx)

**Supplementary Table 3. ISS Probe panel**

List of RNA probes with gene name and ENSEMBL ID for each gene. Probes were obtained through CARTANA. (See SuppTable_3.xlsx)

**Supplementary Table 4. Enriched pathways across osteogenesis pseudotime**

Pathways enrichment scored from gene-sets obtained through numerous databases against pseudotime-associated genes within the osteogenesis subcompartment. (See SuppTable_4.xlsx)

**Supplementary Table 5. Craniosynostosis-associated genes**

Curated list of craniosynostosis genes obtained from the online platform Genomics England Panel. (See SuppTable_5.xlsx)

**Supplementary Table 6. Drugs with teratogenicity warning**

List of drug names obtained through chEMBL database with black box labels of teratogenicity as a feature. (See SuppTable_6.xlsx)

**Supplementary Table 7. RNAscope probes**

List of probes used for RNAscope experiments and relevant information (See SuppTable_7.xlsx)

**Supplementary Table 8. Genome-wide association studies metadata**

Details of GWAS studies utilised in fGWAS enrichment analysis (See SuppTable_8.xlsx)

**Supplementary Table 9. SCENIC+ results**

TF-enhancer-gene links for osteogenesis, chondrogenesis, fibrogenesis, early joint progenitors, immune and Schwann cells. (See SuppTable_9.xlsx)
