## Extended Data Figures for "A multiomic atlas of human early skeletal development"

### Extended data figures for:
### A multiomic atlas of human early skeletal development

[1]Ken To*, [1]Lijiang Fei*, [1]J. Patrick Pett*, [1]Kenny Roberts, [1]Krzysztof Polański, [1]Tong Li, [1,2]Nadav Yayon, [1,2]Peng He, [1]Chuan Xu, [1]James Cranley, [1]Ruoyan Li, [1]Kazumasa Kanemaru, [1]Ni Huang, [1]Stathis Megas, [1]Laura Richardson, [1]Rakesh Kapuge, [1]Shani Perera, [1]Elizabeth Tuck, [1]Anna Wilbrey-Clark, [1]Ilaria Mulas, [1]Fani Memi, [1]Batuhan Cakir, [1]Alexander V. Predeus, [1,3]David Horsfall, [1]Simon Murray, [1]Martin Prete, [1]Pavel Mazin, [4]Xiaoling He, [1]Kerstin B. Meyer, [1,3]Muzlifah Haniffa, [4,5]Roger A. Barker, [1]Omer Bayraktar, [6]Christopher D. Buckley, [1#]Sarah A. Teichmann

*These authors contributed equally
[#]corresponding author

[1] Wellcome Sanger Institute, Wellcome Genome Campus, Hinxton, UK

[2] European Molecular Biology Laboratory, European Bioinformatics Institute (EMBL-EBI), Wellcome Genome Campus, Cambridge, UK

[3] Newcastle University, Biosciences Institute, Newcastle University, Newcastle upon Tyne, UK

[4] John van Geest Centre for Brain Repair, Department of Clinical Neurosciences, University of Cambridge, Cambridge, UK

[5] Wellcome-MRC Cambridge Stem Cell Institute, University of Cambridge, Cambridge, UK

[6] Kennedy Institute of Rheumatology, University of Oxford, Oxford, UK

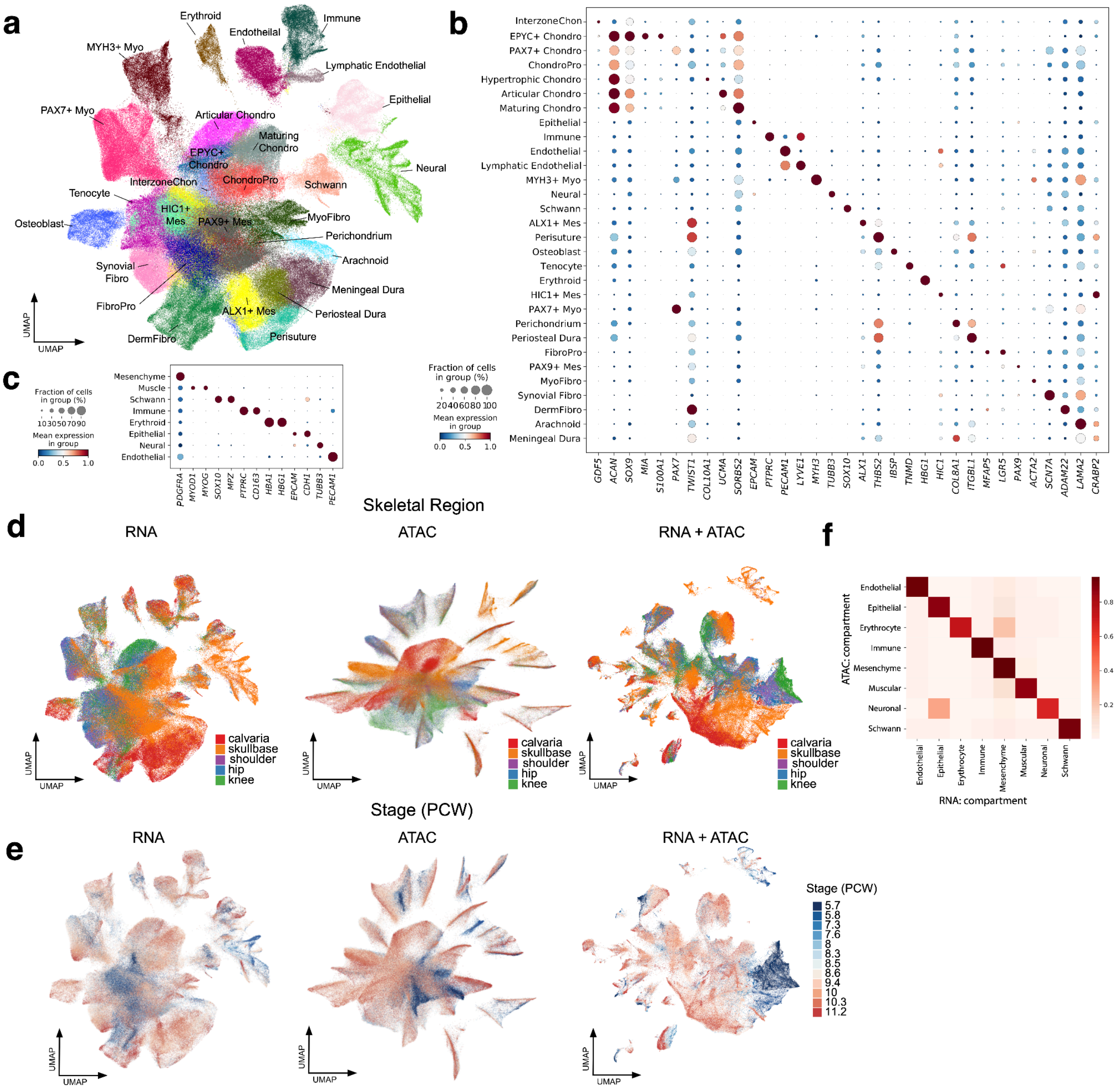


**Extended Data Fig. 1 Droplet dataset overview**

**a.** UMAP-embedding of RNA droplets with broad cell cluster labels **b.** Dotplot with marker genes per broad cluster **c.** Dotplot with marker genes for each cell type compartment **d.** Skeletal region metadata displayed on RNA (left), ATAC (middle) and co-embedding (Right). **e.** Stage (PCW) metadata displayed on RNA (left), ATAC (middle) and co-embedding (Right). **f.** Heatmap of concordance of broad cell compartment labels across RNA and ATAC modality, scale bar shows the number of intersected cells of RNA and ATAC compartment (scaled by row).

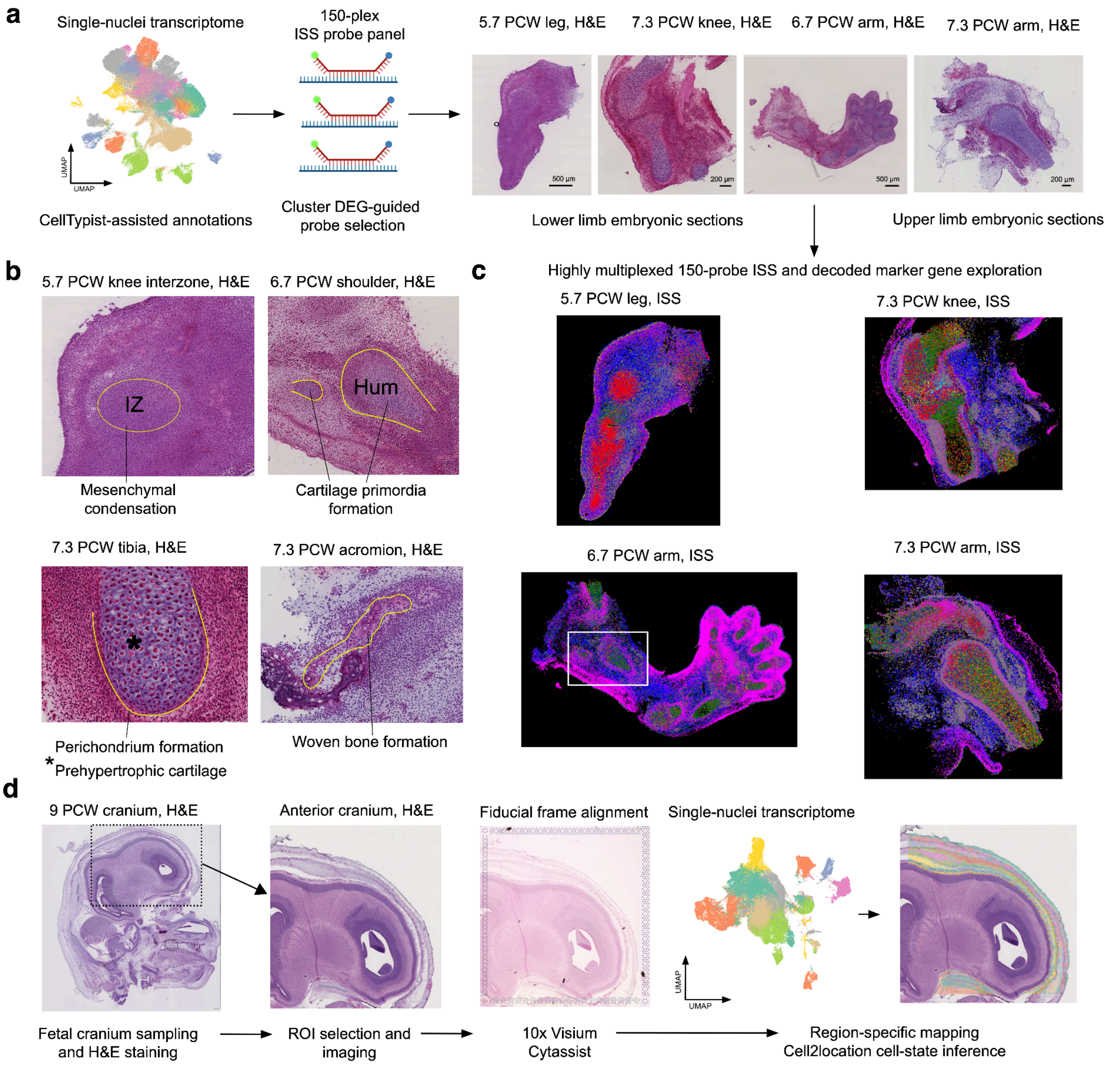


**Extended Data Fig. 2 Spatial transcriptomics analysis and workflow**

**a.** UMAP-embedding of RNA droplets of the appendicular joints manually annotated informed by Celltypist utilising Zhang et al, 2023 labels. Per-cluster DEG used to design 150-plex probe panel, subsequently applied to sectioned samples of 5-7 PCW synovial joints (Right). **b.** Regions of cellular niches within the samples utilised for ISS imaging. **c.** Visualisation of probe distribution on ISS imaging data applied to histological sections in a, white box shows nascent shoulder joint in 6.7 PCW arm. **d.** Workflow for 10x Visium Cytassist applied to 9 PCW cranium.

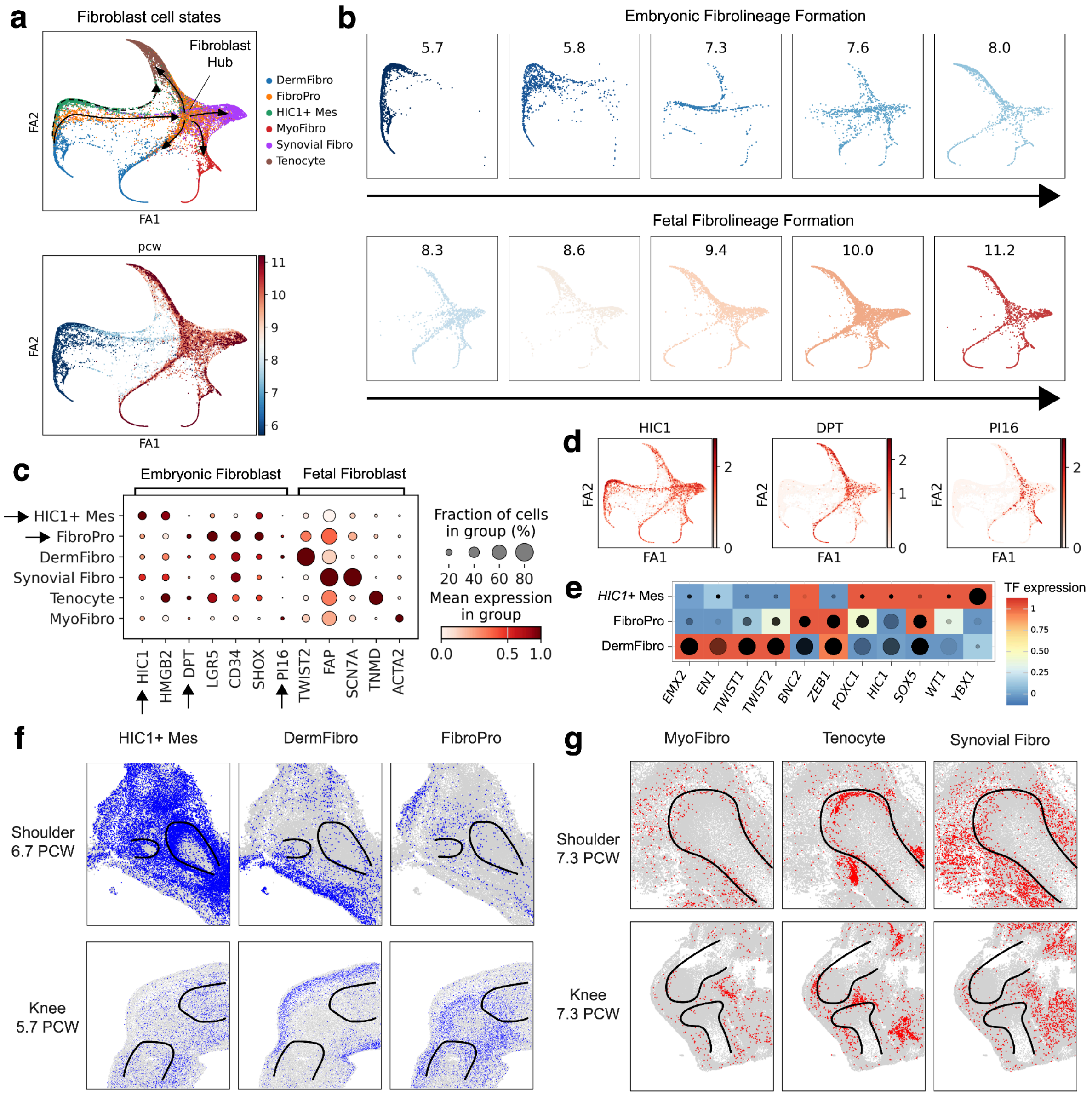


**Extended Data Fig. 3 Appendicular joints fibroblast cell states**

**a.** Force-directed embedding of Fibroblast cell states. Dotted arrow showing tenogenesis trajectory, solid arrows showing fibroblast formation trajectory, leading to formation of fibroblast hub at fetal stages (Top). **b.** Stage (PCW) metadata displayed over force embedding, scale bar shows PCW. **c.** Dotplot with marker genes per cell state. Solid arrows show markers of comparable clusterss described in the mouse (*HIC1*$^{+}$ mesenchyme, and *PI16*+*DPT*+ Universal fibroblasts). **d.** Visualisation of marker genes on embedding. **e.** TF expression across three fibroblast clusters. Color shows normalised expression, dot size shows target gene accessibility (AUCell) and dot shade (grayscale) shows target gene expression (GEX AUCell). **f.** Imputed fibroblast cell states in the embryonic limb < 7 PCW. **g.** Imputed fibroblast cell states in the fetal limb > 7 PCW.

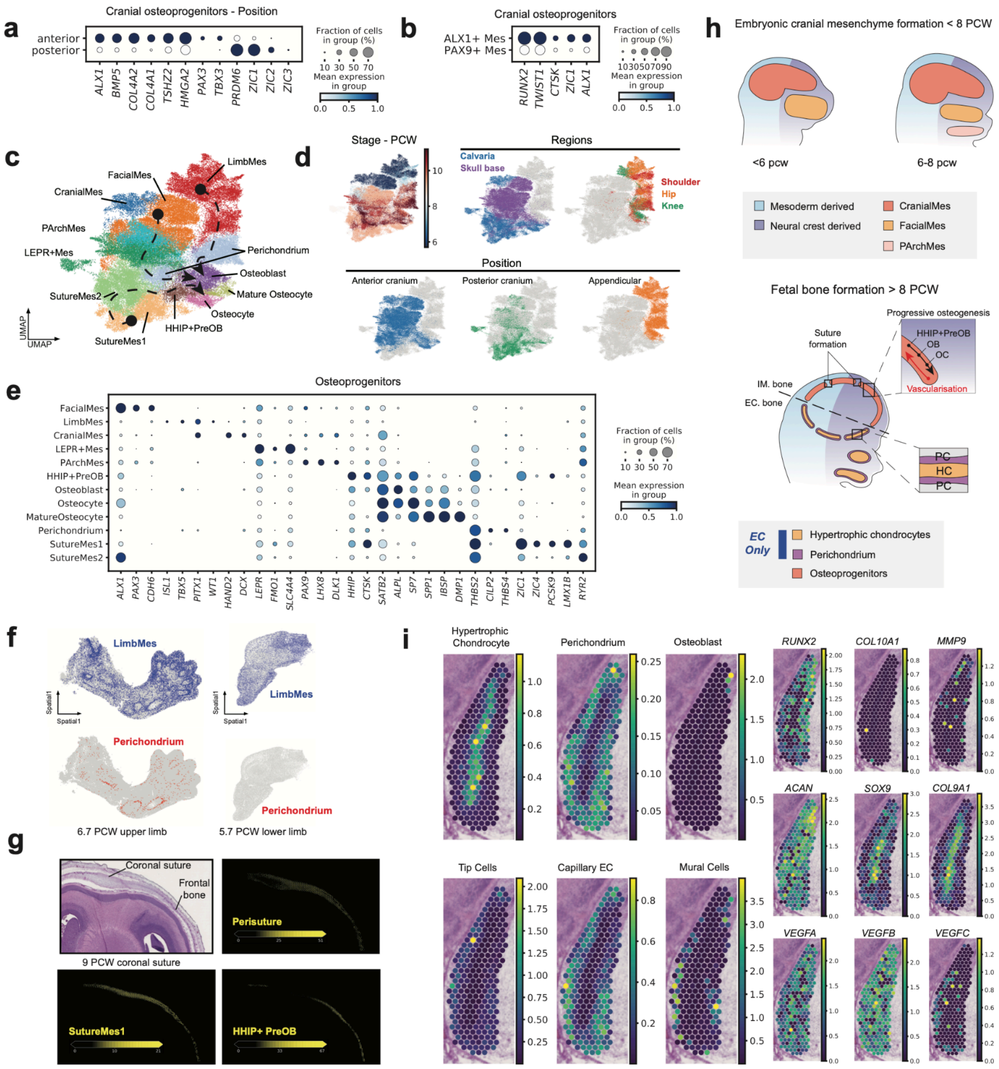


**Extended Data Fig. 4 Osteogenesis cell states**

**a.** Dotplot with marker genes per relative anatomical position in the cranial osteogenic cell states. **b.** Dotplot with marker genes of previously described mouse suture cells across broad osteogenic clusters. **c.** UMAP embedding of osteogenesis sub-compartment. **d.** Stage (PCW), region and position metadata overlaid on UMAP embedding. **e.** Dotplot of marker genes per subcompartment osteogenic cluster. **f.** Imputed osteogenic cell states in the appendicular ISS data **g.** Cell2location cell state enrichment in the coronal suture and frontal bone **h.** Schematic showing cranial progenitor maturation across embryonic (top) and fetal (bottom) stages. **i.** Cell2location cell state enrichment in the sphenoid bone of the skull base (endochondral niche) (Left), gene expression in the same region (right).

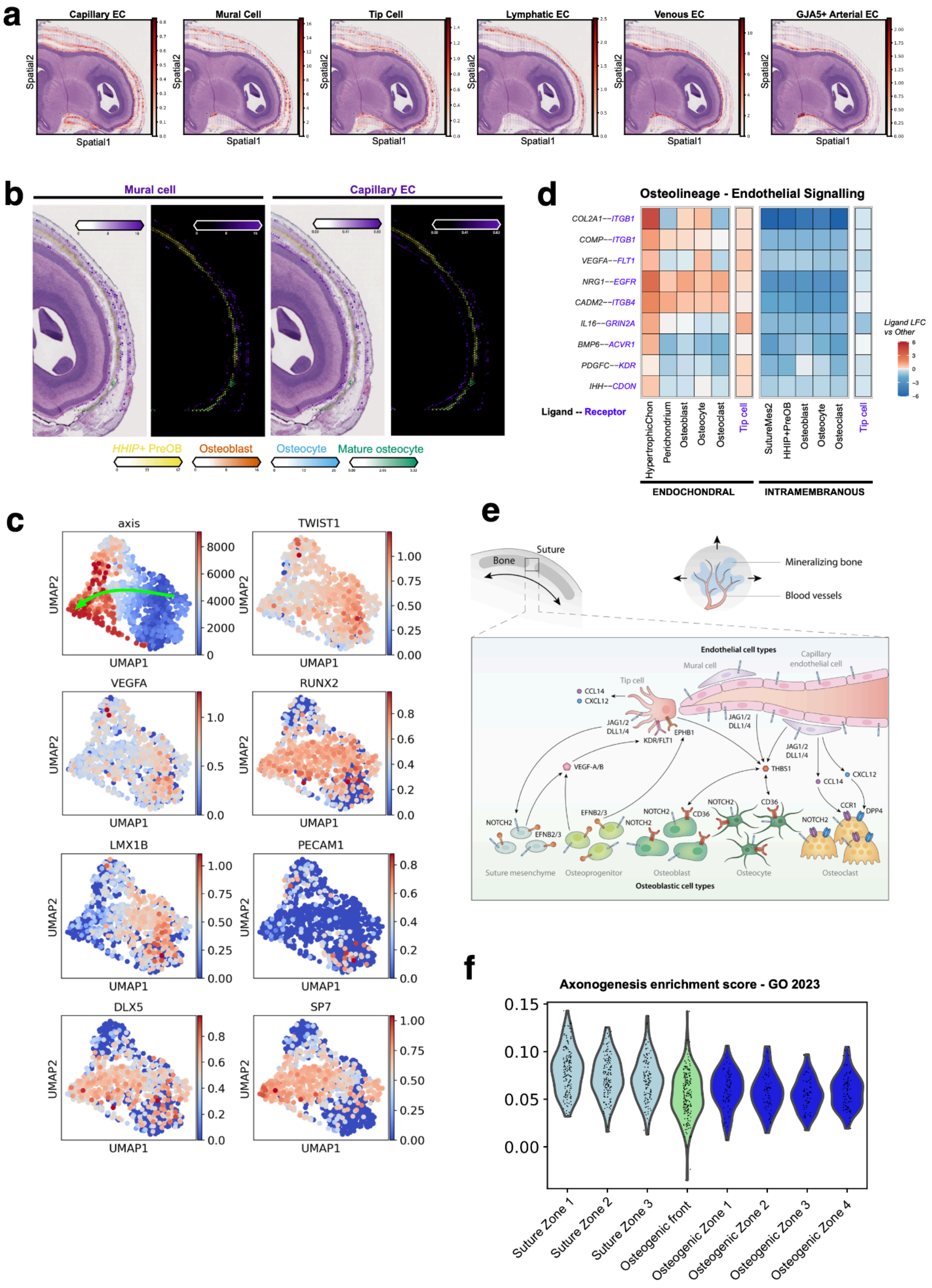


**Extended Data Fig. 5 Cellular interactions within the suture niche**

**a.** Cell2location endothelial cell state enrichment in the anterior portion of the cranium **b.**

Cell2location enrichment of osteogenic cell states alongside mural (left) and capillary EC (right) on the anterior portion of the frontal bone. **c.** Clustering of visium voxels with organ-axis values and gene expression. **d.** NicheNet inferred differential cell-cell interactions between osteogenic cell states and Tip cells in the EC and IM niches. **e.** Schematic of endothelial and osteogenic cell state interactions at the boundaries of the suture mesenchyme. **f.** Enrichment of axonogenesis gene set from the Gene-ontology 2023 database across the organ-axis spatial bins.

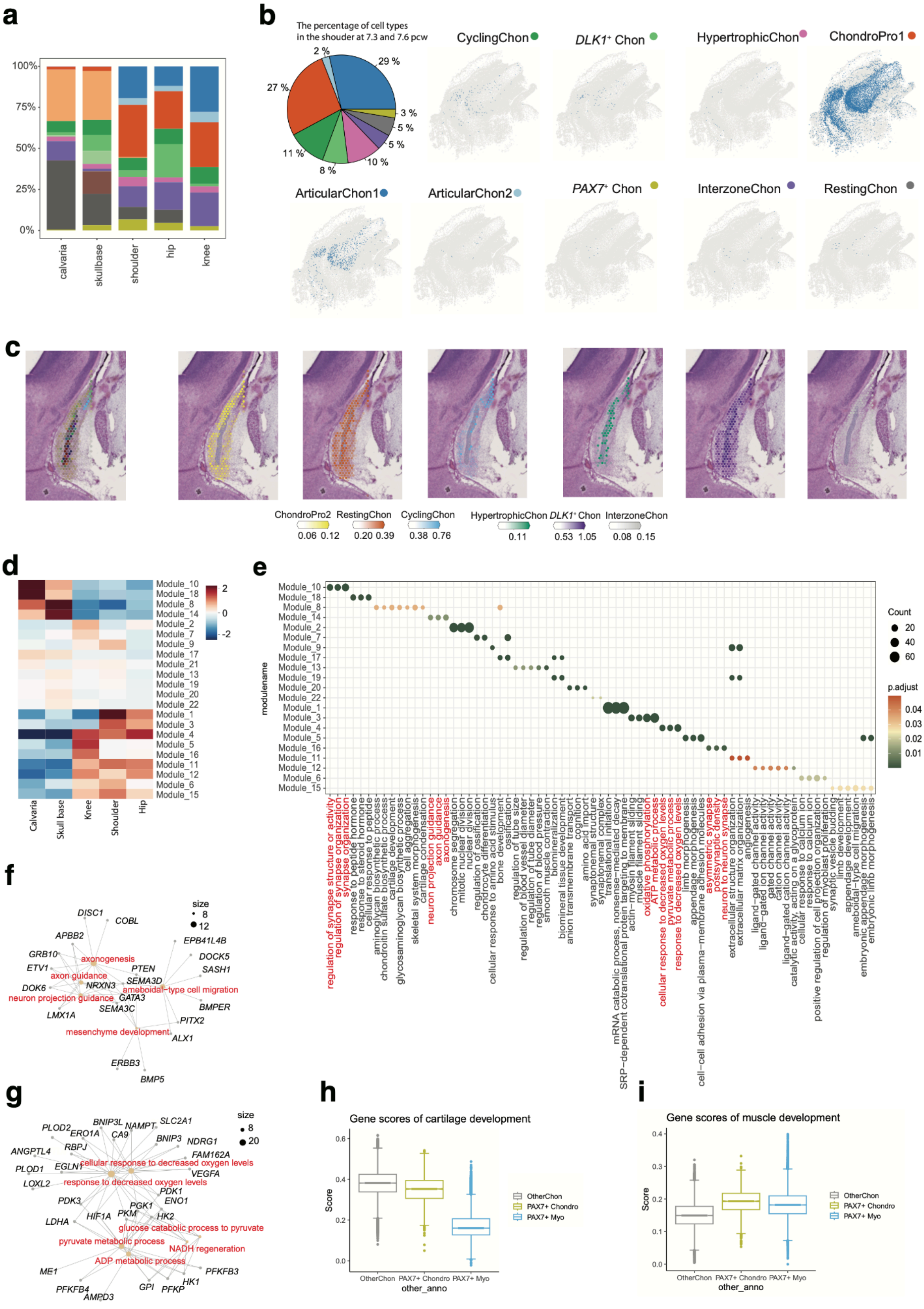

**Extended Data Fig. 6 Chondrogenesis cell states**

**a**. Boxplot showing the distribution of cell cluster abundance of chondrocytes per anatomical region and per PCW. **b**. The left pie chart showing the percentage of cell clusters in the shoulder at 7.3 and 7.6 PCW. The right spatial plots showing the predicted cell clusters from ISS-Patcher cell cluster imputation. Colors represent the corresponding cell clusters. **C**. Histological view and annotations of chondrocytes in the sphenoid sections from 10x Cytassist visium. **d**. Heatmap showing the kernel values of module genes per anatomical region. **e**. Top 3 enriched biological process GO terms for each module. The GO terms are selected based on the gene number and terms of the same gene number remain. Colour represents the p adjusted values, while the dot size represents the gene number of each GO term. **f**. Network visualisations of enriched genes of calvaria-specific GO terms. **g**. Network visualisations of enriched genes of appendicular-specific GO terms. **h**. Boxplot showing the gene scores of cartilage development. Boxplot center line, median; boxes, first and third quartiles of the distribution; whiskers, highest and lowest data points within 1.5 × interquartile ratio (IQR). OtherChon represents the whole chondro-lineages except *PAX7*$^{+}$ Chon. **i**. Boxplot showing the gene scores of muscle development. Boxplot center line, median; boxes, first and third quartiles of the distribution; whiskers, highest and lowest data points within 1.5 × IQR.

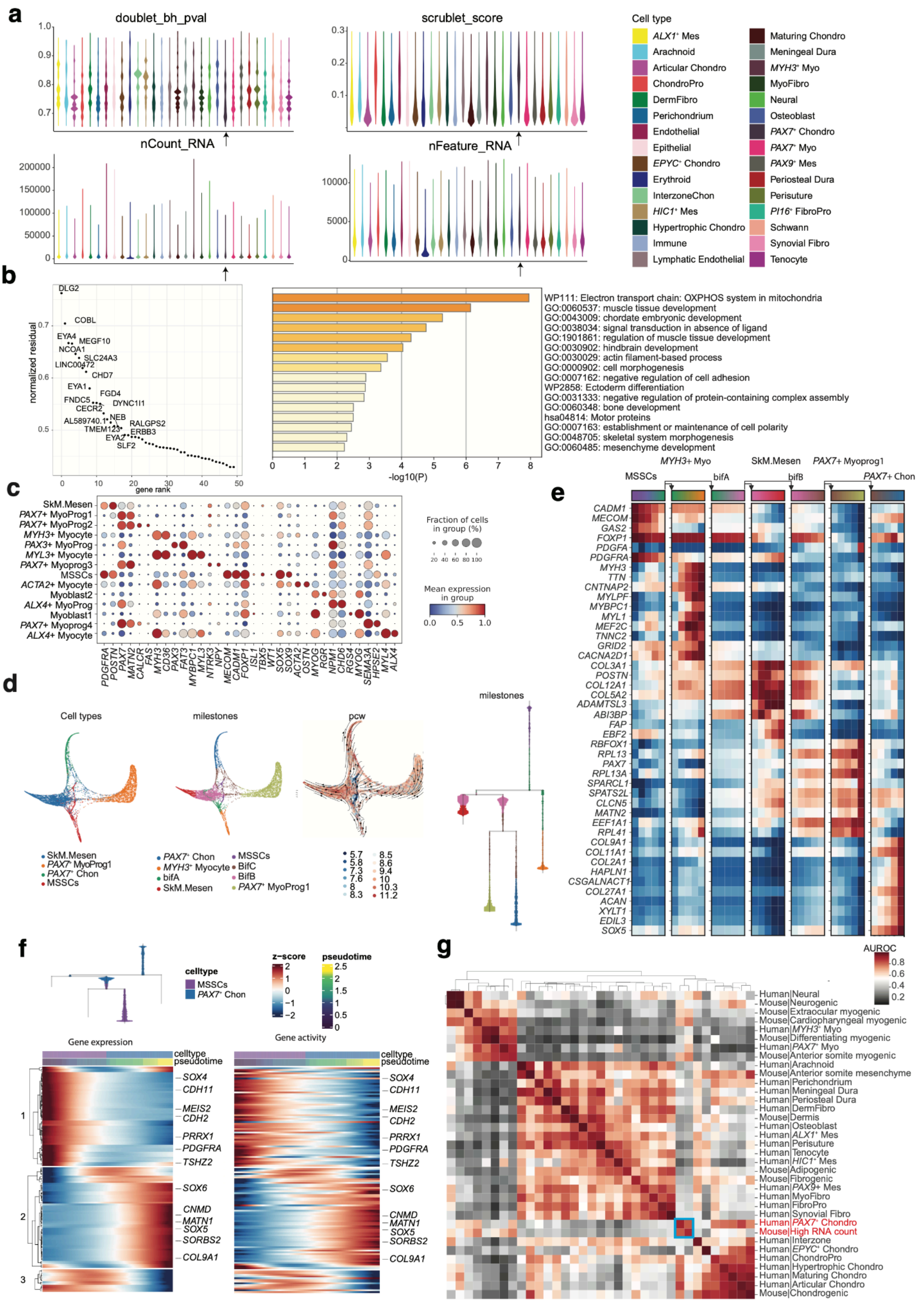

**Extended Data Fig. 7 *PAX7*⁺ Chondrocyte**

**a**. Violin plots showing the adjusted doublet p values, scrublet score, gene count, and gene numbers of each broad cell cluster. The adjusted doublet p values were calculated according to the two-step doublet score processing steps[114]. Black arrow represents *PAX7*$^+$ Chon **b**. Deviation of genes of *PAX7*$^+$ Chon from the linear mix for expression profiles of ChondroPro1, *PAX7*+ Myoprogenitor1 and SkM.Mesen[115]. Barplot showing the enriched GO terms of top50 genes. **c**. Dotplot showing the represented genes of myogenic cell clusters. The dot size represents the fraction of cells in each cell cluster and the dot colour represents the scaled mean expression. **d**. FA layout showing the trajectories and velocity analysis of cell clusters that share the mutations with *PAX7*$^+$ Chon. Milestones (cell states) are inferred from scFate. Dendrogram showing the inferred trajectory tree of milestones. **e**. Heatmap showing the top differentiated expressed genes for each milostone. **f**. Subsetted tree from embryonic skeletal progenitors to *PAX7*$^+$ Chon (Top). Dynamic changes in gene expression and gene activities from embryonic skeletal progenitors to *PAX7*$^+$ Chon along the pseudotime (Bottom). **g**. Heatmap showing the AUROC scores between human (this study) and mouse cell clusters[87] based on the highly variable gene set using MetaNeighbor[116].

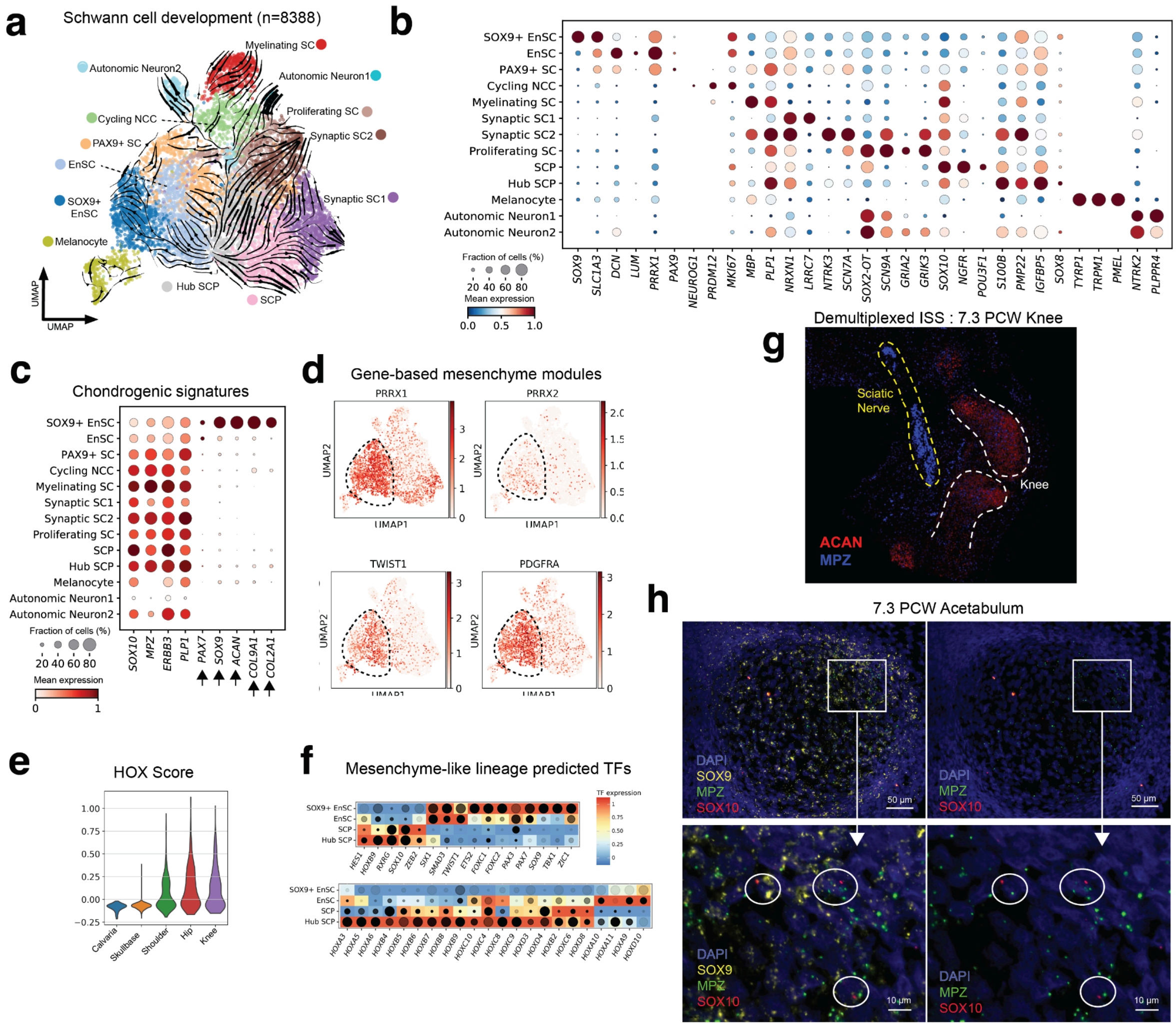

**Extended Data Fig. 8 Schwann cell lineage**

**a.** RNA-velocity arrows displayed on embedding of Schwann lineage cell states, with numerous cell states emerging from Hub SCP (Schwann cell precursor). **b.** Dotplot showing marker gene expression across Schwann cell states. **c.** Dotplot showing chondrogenesis-associate genes across Schwann cell states. Black arrows indicate distinct markers of the *SOX9*+ enSC population. **d.** Expression of mesenchyme associated genes in the Schwann UMAP. **e.** Enrichment of HOX genes per anatomical region of the Schwann compartment **f.** TF expression across the putative chondrogenic lineage of the Schwann clusters. Color shows normalised expression, dot size shows target gene accessibility (AUCell) and dot color (grayscale) shows target gene expression (GEX AUCell). **g.** Marker genes of ISS RNA probes in the ISS image of the knee joint demonstrating nerve-associate enrichment of Schwann markers, and the presence of *MPZ* staining in the developing bone. **h.** Three-plex RNAscope of marker genes in the acetabulum demonstrating enrichment and colocalization of markers of SOX9$^{+}$ enSC.

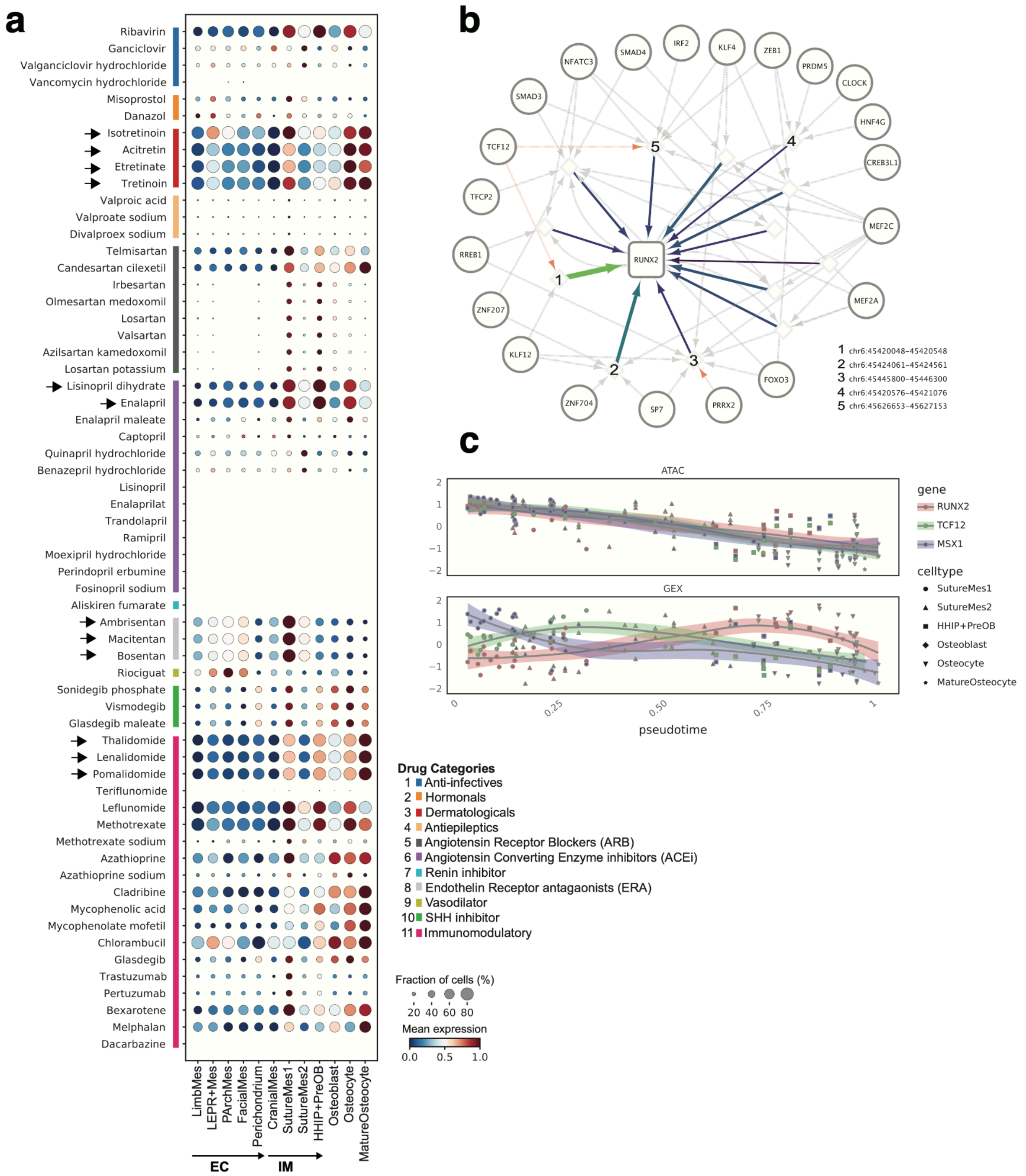


**Extended Data Fig. 9 Drug2Cell analysis and osteogenic network**

**a.** Dotplot with drug2cell (chEMBL) enrichment of teratogenic drugs across cell states of the osteogenic trajectory. Selected highly enriched drugs are marked with arrows (y-axis) and cell states unique to either endochondral (EC) or intramembranous (IM) ossification are marked on the x-axis. Patterns of different drugs predicted to affect target genes at different parts of the trajectory are visible. **b.** eGRN showing the regulation of RUNX2 predicted with SCENIC+. Various TFs (circles) are predicted to regulate RUNX2 (square) via binding to regulatory elements (diamonds). The colour and width of arrows from regions to RUNX2 show importance scores. Links from two TFs appearing in Fig. 2 and 6 are highlighted in red.

Various osteogenic TFs, including SP7, are predicted to regulate RUNX2. **c.** Accessibility and expression of three selected genes along pseudotime of the osteogenic trajectory (Fig. 3). Higher expression of inhibitory TFs is observed at the beginning (SutureMes1/2), while RUNX2 shows higher expression towards the end of the trajectory. Similar accessibilities suggest additional layers of or non-pioneering factor driven regulation.
